# Proteome-wide base editor screens to assess phosphorylation site functionality in high-throughput

**DOI:** 10.1101/2023.11.11.566649

**Authors:** Patrick H Kennedy, Amin Alborzian Deh Sheikh, Matthew Balakar, Alexander C Jones, Meagan E Olive, Mudra Hegde, Maria I Matias, Natan Pirete, Rajan Burt, Jonathan Levy, Tamia Little, Patrick G Hogan, David R Liu, John G Doench, Alexandra C Newton, Rachel A. Gottschalk, Carl de Boer, Suzie Alarcón, Gregory Newby, Samuel A. Myers

## Abstract

Signaling pathways that drive gene expression are typically depicted as having a dozen or so landmark phosphorylation and transcriptional events. In reality, thousands of dynamic post-translational modifications (PTMs) orchestrate nearly every cellular function, and we lack technologies to find causal links between these vast biochemical pathways and genetic circuits at scale. Here, we describe “signaling-to-transcription network” mapping through the development of PTM-centric base editing coupled to phenotypic screens, directed by temporally-resolved phosphoproteomics. Using T cell activation as a model, we observe hundreds of unstudied phosphorylation sites that modulate NFAT transcriptional activity. We identify the phosphorylation-mediated nuclear localization of the phosphatase PHLPP1 which promotes NFAT but inhibits NFκB activity. We also find that specific phosphosite mutants can alter gene expression in subtle yet distinct patterns, demonstrating the potential for fine-tuning transcriptional responses. Overall, base editor screening of PTM sites provides a powerful platform to dissect PTM function within signaling pathways.

## Introduction

Nearly every eukaryotic cellular process is controlled by post-translational modifications (PTMs), which can modulate protein subcellular localization, protein-biomolecular interactions, enzymatic activity, stability, etc. Protein phosphorylation is arguably the best characterized PTM^1^. The human genome encodes for roughly 500 protein kinases and 200 phosphatases that control the coupling and hydrolysis of phosphates, respectively, on substrates in a rapid and dynamic fashion^2,3^. These signaling cascades organize into elaborate biochemical networks that allow the cell to process information about its intra- and extracellular environmental changes. Mass spectrometry-based proteomics has revolutionized our ability to map global signaling pathways at the phosphorylation site (phosphosite)-specific level, across time and cellular space. Current phosphoproteomics experiments can quantify tens of thousands of phosphosites, tracking their dynamics upon cell stimulation, drug treatment, or mutational status^4–10^. Unfortunately, fewer than 3% of the nearly quarter million phosphorylation sites identified have an ascribed function^11,12^. Since functionally characterizing novel modification sites is laborious and resource intensive, we often resort to describing complex biological systems by the limited number of well-characterized phosphosites for which we have good reagents.

Functional genomics has greatly increased our throughput for associating genes with specific cellular phenotypes. Genome-wide CRISPR/Cas9 technology coupled with phenotypic screens allow researchers to identify which genes or non-coding regions are important for a specific function such as gene expression^13^, cytokine secretion^14^, cell proliferation^15^, or cell survival^16–18^. More recently, CRISPR/Cas9-mediated base editors, which introduce specific nucleotide substitutions in genomic DNA rather than double stranded DNA breaks^19^, have been used for mutational scanning across protein coding genes and regulatory elements^20–22^. Base editor technology holds immense promise to study PTM site function in high-throughput by mutating specific amino acids, bypassing the need to create site-specific homology-directed repair templates^23^.

Here, we describe an experimental workflow to study phosphorylation site functionality in high-throughput. By coupling quantitative phosphoproteomics with “proteome-wide” base editing of individual phosphosites and phenotypic screens, we are able to functionally evaluate a large number of previously unstudied phosphosites that are involved in cell proliferation or the transcriptional responses following T cell activation. Applying this “signaling-to-transcription network mapping” to T cell activation, we show that we can recapitulate many known aspects of the pathway, while discovering novel kinase activities and specific phosphorylation events that control different aspects of the transcriptional response. This allowed us to identify a specific phosphosite on the phosphatase PHLPP1 as a novel regulator of T cell activation-induced NFAT and NFκB activities. Transcriptional profiling of PHLPP1 phosphosite mutant T cells shows that individual phosphorylation events differentially impact downstream gene expression in subtle yet distinct patterns, creating the potential for fine-tuned control of gene expression through signaling pathway modulation. PTM-centric base editor screens provide an experimental framework to functionally interrogate and systematically decode the vast network of biochemical signaling events to their downstream phenotypes.

## Results

### Optimization of base editing for experimentally-derived phosphorylation sites

To develop a system by which we could profile and then functionally assess signaling pathways and their effects on gene expression, we focused on a classic model of T cell activation in the human T cell leukemia line Jurkat E6-1, during which multiple kinases and downstream transcription factors are activated (**Figure 1A**)^24^. We performed a temporally-resolved quantitative phosphoproteomic experiment, assaying global phosphorylation patterns for 0, 3, 9 and 27 minutes of T cell activation using α-CD3 and α-CD28 agonist antibodies^25,26^. Of the 26,037 quantified phosphopeptides, 899 were significantly differentially regulated (moderated F test, FDR < 0.05) during this time series (**Figure 1B, Supp Table 1, and Supp Data File 1**). PTM-SEA^27^, a PTM site-centric analog to GSEA^28^ (gene set enrichment analysis), showed that various kinase activities were temporally regulated during the first 30 minutes of T cell activation (**Figure 1C**). This analysis culminated with the perturbation signatures for “anti-CD3” and “phorbol esters”, indicating the temporally-regulated phosphopeptides reflected the appropriate T cell activation pathways.

**Figure 1.**
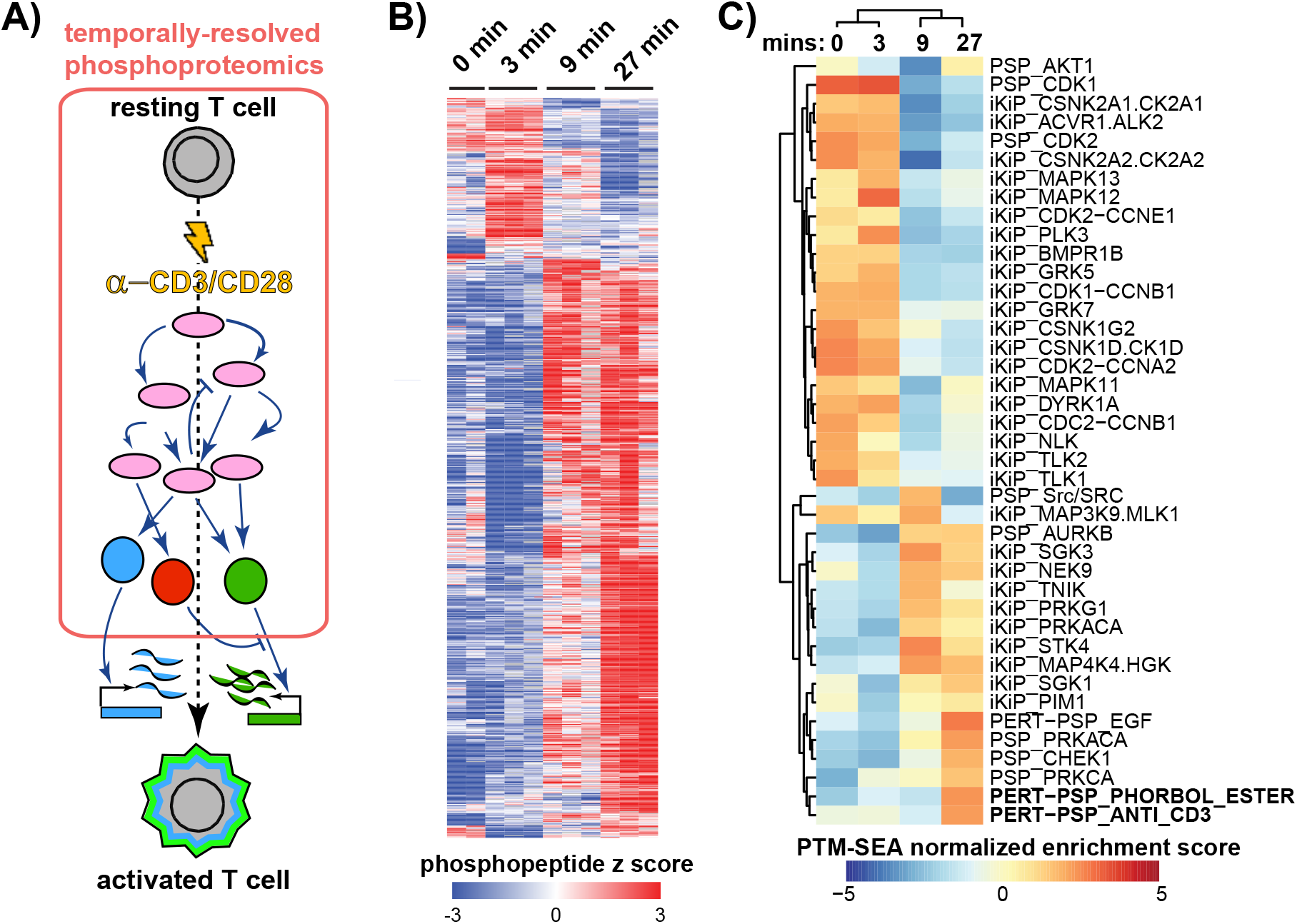
Signaling dynamics of early T cell activation. **A)** Diagram of early T cell activation and where the phosphoproteomics typically samples. Pink ovals represent kinases, colored circles are transcription factors. **B)** Heatmap of statistically regulated phosphopeptides during the first 27 minutes of T cell stimulation. Interactive data is associated with this figure (**Supp File 1**). **C)** Heatmap of PTM-SEA terms for the phosphoproteomics time series data. “iKiP” indicates the *In Vitro Kinase-to-Phosphosite Database*^87^, “pKS” is kinase-substrate pairs from PhosphositePlus (PSP). “PERT” is a PSP-curated perturbation.

Using a custom bioinformatics pipeline, we queried which of the ∼19,000 confidently localized phosphosites we could target using SpCas9-mediated C-to-T or A-to-G editors. We included all detected phosphosites, rather than only temporally regulated ones since there are likely to be phosphosites that were not statistically significant but still contribute to T cell activation. Considering editing windows and targetable locations, we found 7,618 unique phosphosites were targetable with 9,207 distinct sgRNAs using the SpCas9-A-to-G editor ABE8e, while 7,063 unique phosphosites could be targeted by the SpCas9-C-to-T editor BE4 (**Figure 2A and Supp Table 2**)^29,30^. Roughly half of the editable phosphosites overlapped between the two base editors. The amino acid side chain representation of targetable phosphosites reflected those detected and statistically regulated (**Figure 2B**). ABE8e appears to make more structurally conservative missense mutations, and unlike BE4, can target tyrosine-encoding codons (**Figure 2A**).

**Figure 2.**
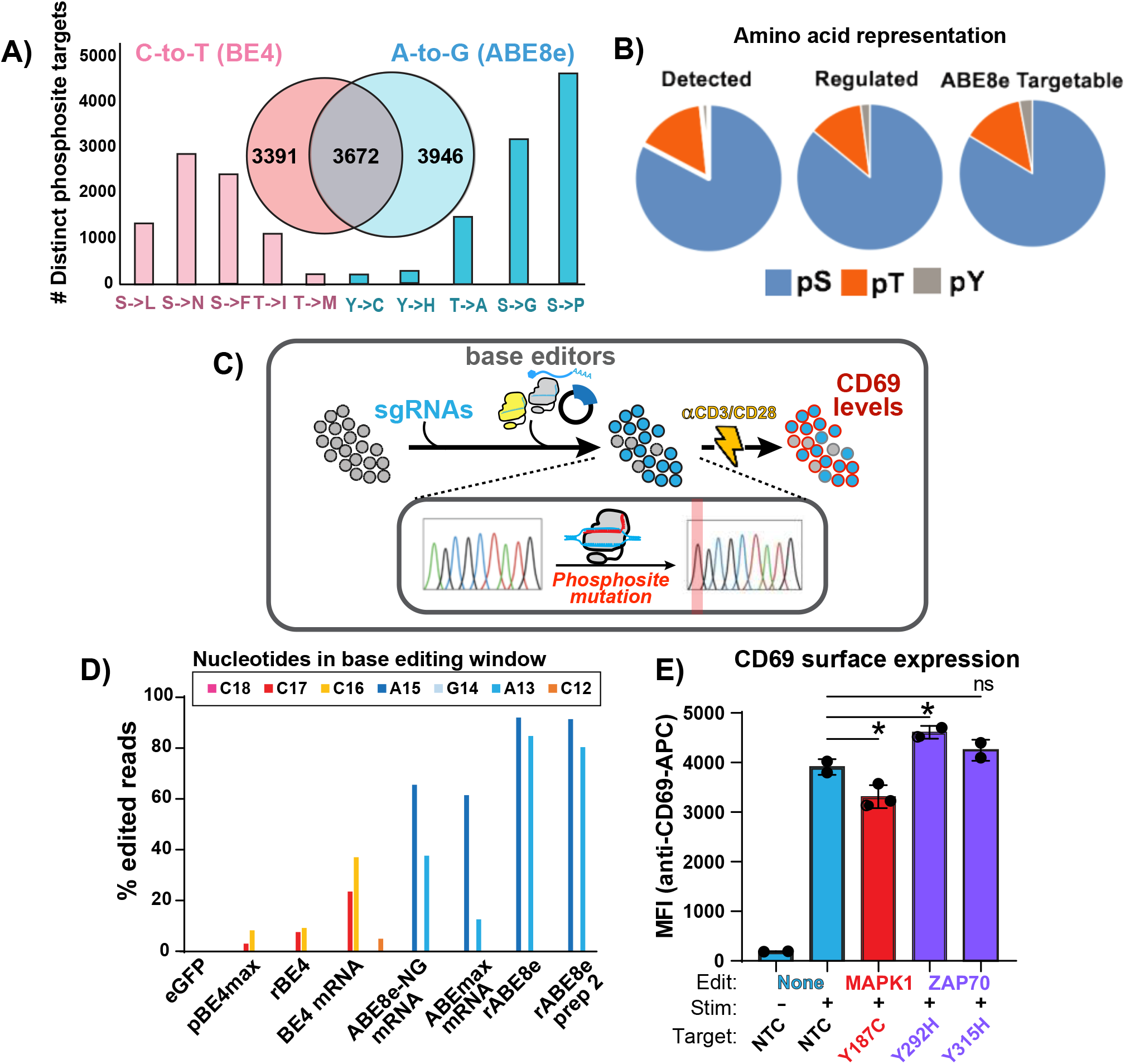
Base editing capabilities of empirically derived phosphorylation sites. **A)** The number of distinct phosphosites targeted by at least one sgRNA and what the resultant amino acid codon will be for BE4 (pink) or ABE8e (blue). The overlap of unique phosphosite targets is in the inset. BE4 cannot mutate Y to any other amino acid and was omitted. **B)** The distribution of phosphorylated amino acid side chains in all phosphopeptides detected by mass spectrometry, those statistically regulated, and the subset that can be targeted by ABE8e. **C)** Diagram for testing various base editor delivery methods in an arrayed format, one sgRNA at a time followed by T cell activation. **D)** Base editing efficiency, as determined by NGS amplicon sequencing in percent of edited reads, testing the nucleofection of different biomolecules to deliver base editors. The base editing window, counting right to left from the PAM sequence, is shown in the inset and the nucleic acid targets are color coordinated. “p” indicates plasmid, “r” is recombinantly expressed and purified, “mRNA” is synthetic, capped mRNA. **E)** Effect of phosphosite base edits on T cell activation-induced CD69 surface levels. “Edits” indicates the targeted gene, “Stim” indicates 12 hours of treatment with α-CD3/CD28 agonist antibodies, and “Target” indicates amino acid targeted. NTC, non-targeting control at the *HEK3* locus; MFI, mean fluorescence intensity. * denoted p-value < 0.05 and “ns” for not significant.

To develop a flexible genomic engineering approach, we based our base editing strategy on previous genome-wide CRISPR/Cas9 screens in primary T cells where sgRNAs are delivered via lentivirus, followed by electroporation of Cas9 protein^15^. We nucleofected several different types of biomolecules to determine the most efficient base editors (**Figure 2C**). Using Jurkat cells stably expressing an sgRNA targeting the model *HEK3* site in humans, we nucleofected either plasmid DNA, chemically synthesized and capped mRNA, or recombinant protein of different base editor versions. We found that purified recombinant ABE8e protein (NGG PAM,) properly edited over 95% of adenosines in the base editing window (**Figure 2D**). To test the reproducibility of the ABE8e protein, we re-expressed and purified the protein^30^. Again, we found that 92% of the adenosines in the base editing window were mutated to guanosine via Sanger sequencing^31^. These results demonstrate that we can reproducibly achieve sufficiently high base editing efficiency with ABE8e protein for high-throughput screens.

### Base editing phosphosites that promote or inhibit markers of T cell activation

We tested whether mutating phosphosites in proteins known to be involved in the TCR signaling pathway with ABE8e protein would affect markers of T cell activation. We base edited the activating tyrosine of MAPK1 (ERK2) Y187, and two targets in the TCR associated kinase ZAP70^32^ (Y292 and Y315) in Jurkat E6.1 cells. Mutant or control cells were activated with α-CD3/CD28 agonist antibodies for 12 hours, stained for the early activation marker surface CD69 levels and analyzed by flow cytometry. Mutation of an inhibitory phosphotyrosine on ZAP70 Y292H^33,34^ increased CD69 surface expression whereas MAPK1 Y187C showed diminished surface CD69 (**Figure 2E**). ZAP70 Y315H showed no effect, consistent with previous reports^35^. Together these data establish that phosphosite mutations can have positive or negative effects on T cell activation levels.

### Phosphosite-centric functional phenotypic screens using pooled base editing for cell proliferation

To assess base editing efficiency in pooled format, we created a lentiviral library consisting of roughly 10,000 phosphosite-targeting sgRNAs for missense mutations, 250 non-targeting controls, and 250 intergenic controls as negative controls. We also included 250 guides that introduce terminating edits in essential genes via mRNA splice site disruption, effectively knocking out the gene. TPR (triple parameter reporter) Jurkat cells, which have individual fluorescent reporters driven by separate NFAT, NFκB, and AP1 transcriptional response elements^36^, were transduced at a multiplicity of infection of 0.3. After puromycin selection, a 500x library coverage aliquot of cells were collected and the rest were electroporated with ABE8e protein. To confirm our base editing was efficient we analyzed the representation of sgRNAs of the controls, comparing pre- and post-ABE8e protein electroporation (**Figure 3A**). The representation of sgRNAs disrupting splice junctions in 250 essential genes was significantly lower six days post-base editing compared to all sgRNAs in the library (**Figure 3B**) indicating our base editing approach was working efficiently. The relative representation of intergenic and non-targeting controls was not affected by introduction of ABE8e protein.

**Figure 3.**
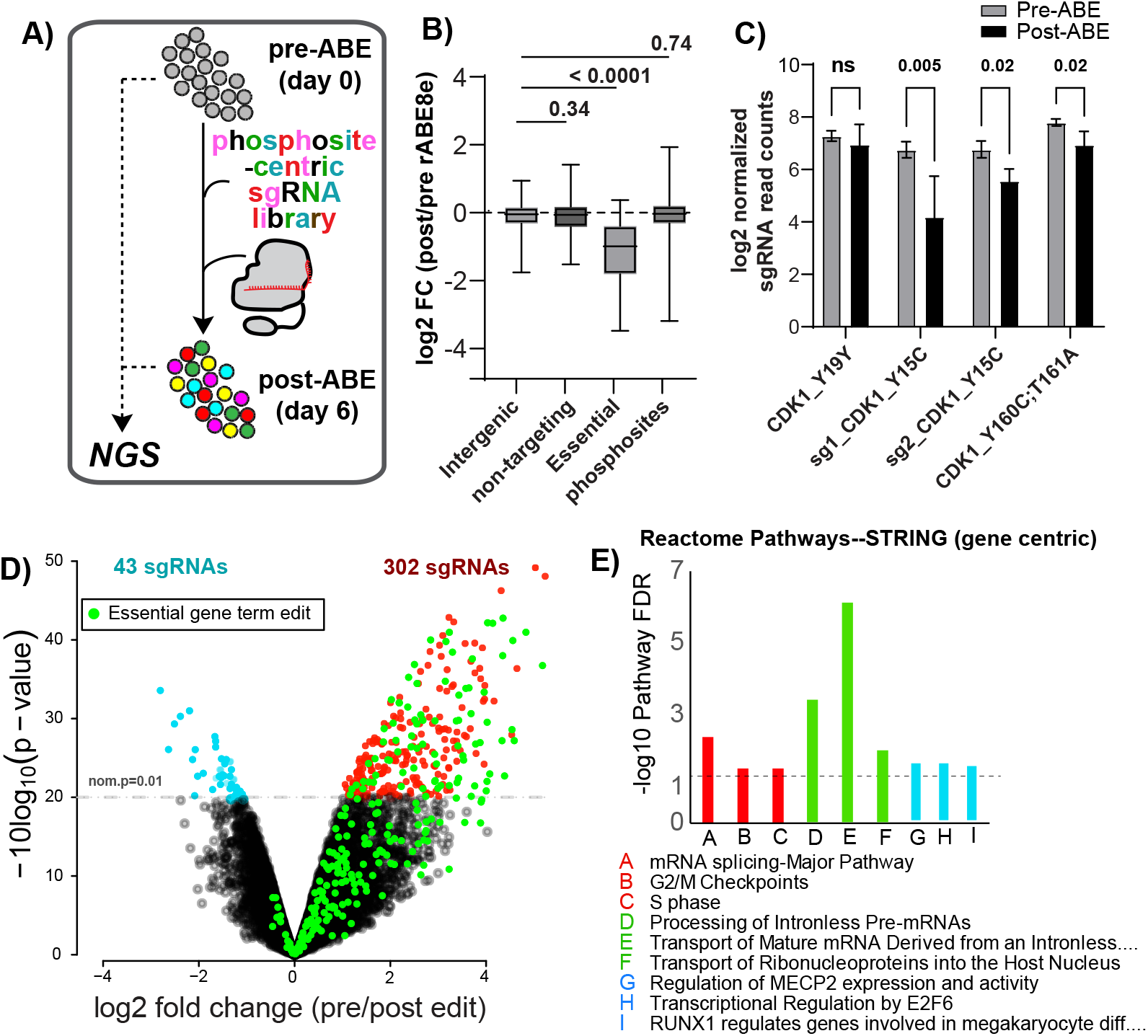
Base editing screening reveals phosphosites involved in proliferation or survival. **A)** Diagram of pooled base editor screens for phosphosites or terminating edits of essential genes important for cell proliferation or survival followed by next-generation sequencing (NGS). **B)** Log2 fold change between pre- and post-ABE8e protein introduction of cells expressing the sgRNA library. “Essential” refers to a terminating edit into essential genes. Intergenic base edits and non-targeting controls are also shown. T test p values are shown. **C)** Mutations made to CDK1 phosphosites and their influence of sgRNA representation before and after base editing in a pooled format. “sg1 and sg2” indicate that two different sgRNAs were used for the Y15C mutation. T-test p values are shown. **D)** Volcano plot showing the distribution of all sgRNAs pre- and post-ABE8e protein introduction. Green points indicate terminating edits in essential genes. Red points indicate statistically significant sgRNA targeting phosphosites depleted after base editing. Blue points are sgRNAs targeting phosphosites that were enriched after base editing. **E)** Gene ontology analysis of the genes targeted by the sgRNAs depicted in C). Colors coordinate with D). Dotted line is the FDR threshold.

We next examined whether mutation of phosphosites important for cell division would affect cell viability or proliferation in our pooled format without specific stimuli or selective pressure^37^. Examination of phosphorylation sites on CDK1, a kinase whose activity is necessary for proper cell division, showed that Y160C;T161A and Y15C reduced cell viability post-ABE8e electroporation similar in magnitude and direction to levels previously seen using homology-directed recombination^38^ (**Figure 3C**). The silent mutation of Y19Y showed no effect. Analysis of the whole phosphosite-mutant dataset resulted in 302 sgRNAs that were significantly enriched pre-ABE compared to post-editing (**Figure 3D and Supp Table 3**). sgRNAs targeting phosphosites that were depleted after ABE8e protein introduction were enriched for genes involved in the term “Cell Cycle” amongst the top three Reactome pathways (**Figure 3E**). 43 sgRNAs introducing phosphosite mutations were enriched post-base editing, suggesting a proliferative advantage (**Figure 3B**). These genes belonged to the Reactome signatures MECP2 activity, E2F6 and RUNX1 transcriptional regulation, pathways involved in the proliferation of glioblastoma and acute myeloid leukemia cells^39–44^ (**Figure 3E**). PTM site-centric pathway analyses can identify groups of phosphosites (in this case, mutations of phosphosites) enriched in the pre- or post-edited pools. Kinase Library^45^, which uses primary sequence motifs derived from biochemical kinase reactions to predict kinase activity through motif enrichment, identified CDK1/4/5/6/13/18 motifs from aggregated phosphosite mutants depleted relative to pre-edited cells (**Supp Fig 2**). MELK1, NIM1, DYRK2/4, and YANK2/3 motifs were enriched in the pre-edited cells. Together, these data show that our approach of base editing phosphorylation sites in a pooled format can identify phosphorylated residues and putative kinases important for proper cell cycle proliferation or survival.

### Coupling functional phosphosite screens with transcriptional reporters identifies novel regulators of NFAT transcriptional activity

To screen for phosphosites that are functionally linked to transcriptional outputs, we performed a “proteome-wide” base editor screen in activated TPR Jurkat cells, utilizing the NFAT-GFP transcriptional reporter. TPR Jurkat cells stably integrated with the sgRNA library described above were electroporated with ABE8e protein, stimulated with α-CD3/CD28 antibodies for 12 hours, and sorted for high and low GFP (NFAT activity) levels (**Figure 4A**). Genomic DNA was collected from the cells sorted into high and low bins, sgRNA identifiers were amplified by PCR, and were sequenced by next-generation sequencing (NGS). Total read-normalized, log transformed sgRNA counts were moderately to strongly correlated indicating sufficient data quality between transduction quadruplicates (**Supp Figure 4A**). We used the Model-based Analysis of Genome-wide CRISPR/Cas9 Knockout (MAGeCK) to identify and rank which phosphosite mutations, the targets of the sgRNAs, regulate the NFAT transcriptional reporter^17^. Individual sgRNA frequencies showed little bias in differential enrichment based on number of read counts (**Supp Figure 4B and Supp Table 4**). Phosphosites with multiple sgRNAs (**Supp Fig. 4C)** were combined in MAGeCK and the high and low GFP bins were compared. We identified 411 sgRNAs enriched in the GFP high bin, and 293 in the GFP low bin (**Figure 4B and Supp Table 4**). Rolling up our phosphosite level perturbations to the gene level (gene-centric), we performed pathway analyses, using multiple tools, to assess the fidelity of our approach. Enrichment analysis using MAGeCKFlute^46^ identified TCR pathways as enriched in the GFP low (**Figure 4C**). LKB1 and ATM pathways were enriched in the GFP high bin (**Figure 4C and Supp 4D**). g:Profiler^47^, a Gene Ontology-based analysis, also identified TCR pathway in the GFP low bin (**Supp Figure 4D**). GSEA^28^ identified the “TCR *Calcium* Pathway” signature in the GFP high bin (**Supp Figure 4E**), likely due, in part, to dephosphorylation of NFAT as a regulatory mechanism^48–51^.

**Figure 4.**
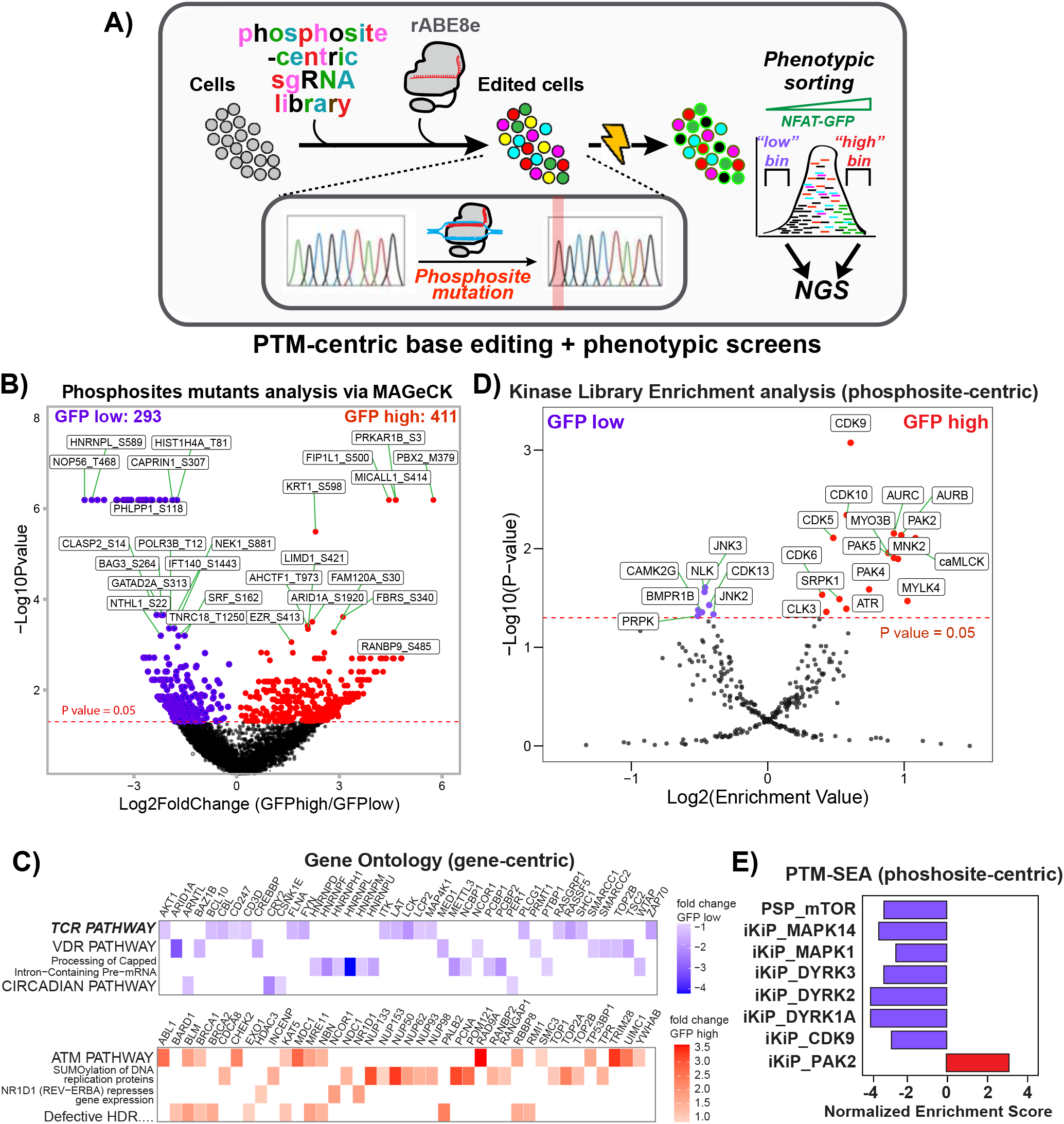
Proteome-wide base editing of phosphosites modulating NFAT transcriptional activity. **A)** Diagram of PTM-centric, proteome-wide base editing coupled to NFAT-GFP transcriptional reporter followed by next-generation sequencing (NGS). **B)** Volcano plot comparing phosphosite edits in GFP high (red) compared to GFP low (purple) bins as determined by MAGeCK. **C)** MAGeCKFlute gene-centric pathway analysis of genes with mutated phosphosites enriched in the GFP low (purple) or GFP high (red) bins. Genes in the respective pathways and their fold change are shown. **D)** Kinase Library, site-centric enrichment analysis of phosphosite mutants enriched in the GFP low (purple) or GFP high (red) bins. Enrichment Values were determined by MAGeCK analysis. **E)** PTM-set Enrichment Analysis (SEA) of phosphosites mutated by ABE8e protein and enriched in the GFP low (purple) or GFP high (red) bins. “iKiP” indicates the *In Vitro Kinase-to-Phosphosite Database*^87^, “pKS” is kinase-substrate pairs from PhosphositePlus (PSP).

Utilizing PTM site-centric analyses, Kinase Library identified several kinases implicated in NFAT activity regulation, such as JNK^52^, NLK^53,54^, and CAMK2G^55^, enriched in the GFP low bin (**Figure 4D**). The GFP high bin was enriched for the CDK9 motif, a kinase involved in general transcriptional regulation^56^, as well as motifs of kinases known to be involved in T cell activation such as PAK2^57^ and CDK5 ^58,59^. CLK3, PAK4, SRPK1 and MYLK4 were also enriched in the GFP high bin but have poorly characterized roles in controlling NFAT transcriptional activity (**Figure 4D**). PTM-SEA^27^ agreed with the Kinase Library results for PAK2 and CDK9 (**Figure 4E**). However, PTM-SEA also identified mutations of MAPK1, MTOR, and DYRK1A/2/3 substrates to be enriched in the GFP low bins, corroborating these kinases’ involvement in T cell activation (**Figure 4E**). As expected, these results demonstrate that phosphosite mutations can directly implicate their involvement in regulating NFAT transcriptional activity, recapitulating known signaling pathways that were constructed from various studies of general T cell activation. These results also strongly suggest that our approach of base editing phosphosites is not only capable of rediscovering crucial signaling molecules in T cell activation, but provides new insights by identifying novel, regulatory kinases and phosphorylation events.

### PTM-centric base editor screening identifies the uncharacterized phosphorylation-mediated nuclear localization of PHLPP1

PHLPP1 is a protein phosphatase most-widely implicated in AKT signaling in cancer^60–62^. In macrophages, PHLPP1 attenuates the JAK/STAT axis by dephosphorylating STAT1^63^. The post-translational mechanisms controlling PHLPP1 function, and its involvement in T cell biology^64^, are poorly understood. We identified the mutation PHLPP1 S118P in our base editing screen to have a strong, negative impact on NFAT transcriptional activity, on par with MAPK1 Y187C (**Figure 5A**). We nucleofected ribonucleoproteins comprised of *in vitro* transcribed sgRNA coupled to ABE8e protein to validate our screen results. The intergenic mutation at the *HEK3* site was used as an editing control, and a terminating edit in *LCP2* or the MAPK1 Y187C mutation abolished or altered T cell activation, respectively, acted as positive perturbation controls. MAPK1 Y187C and PHLPP1 S118P both showed diminished NFAT transcriptional activity (**Figure 5B**). However, unlike MAPK1 Y187C, PHLPP1 S118P showed a small but statistically significant increase in CFP levels (NFκB activity reporter). To extend this analysis, we performed transcriptional profiling of Jurkat T cells harboring the *HEK*3, *LCP2* terminating edit, the MAPK1 Y187C, or the PHLPP1 S118P mutations for zero and six hours post CD3/CD28 stimulation. *LCP2*-terminated cells showed no effect of transcriptional activation six hours after activation, whereas MAPK Y187C showed an intermediate pattern compared to the *HEK*3 controls (**Figure 5C, Supp Table 5, and Supp Data File 2**). The PHLPP1 mutant cells were similar in their transcription patterns compared to the *HEK*3 editing controls, though differences were apparent (**Figure 5C**). We highlighted the genes differentially expressed between *HEK*3, MAPK1 Y187C, and PHLPP1 S118P, and plotted them alongside the *LCP2* terminating edit cells (**Figure 5D**). After k-means clustering and gene ontology analysis of differentially expressed genes, clusters one, five, and six, which were higher in the *Phlpp1* mutant compared to the *HEK3* control identified multiple terms associated with NFκB signaling (NF-kappaB complex, TNFR signaling), corroborating our transcriptional reporter results (**Figure 5B & D**). The MAPK1 Y187C mutant cells were enriched for genes in sterol and isoprenoid biosynthesis (**cluster 4 of Figure 5D**). These results suggest different phosphorylation sites in the T cell activation pathway can regulate downstream gene expression in disparate ways.

**Figure 5.**
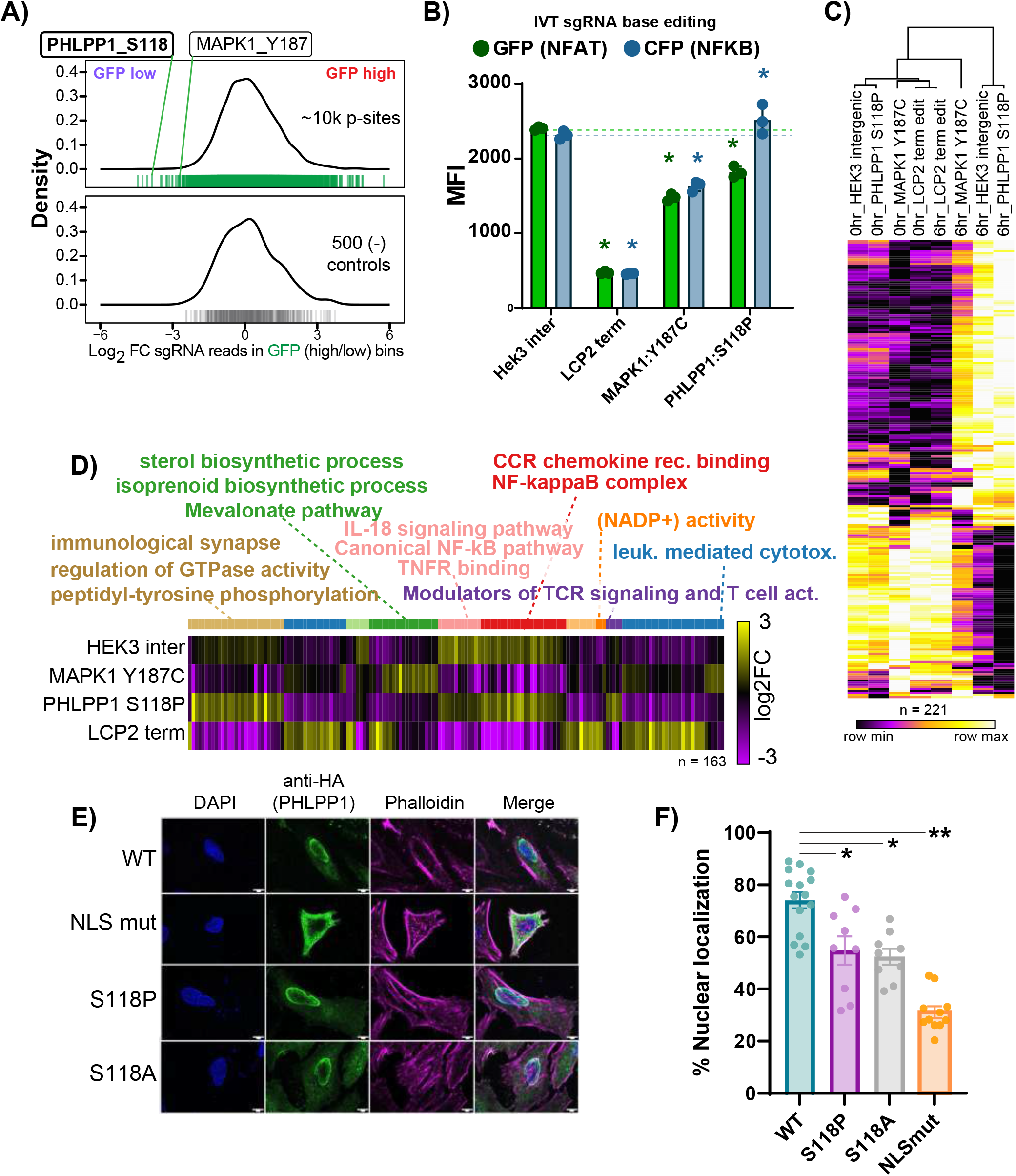
Phosphorylation-induced nuclear translocation of PHLPP1 promotes NFAT and represses NFκB transcriptional responses. **A)** Distribution of log2 fold changes of the ∼10,000 sgRNAs inducing phosphorylation mutations in the NFAT (GFP) transcriptional activity screen (top, green) and non-targeting and intergenic controls (lower panel, gray) between GFP low and GFP high bins. PHLPP1 S118P and MAPK1 Y187C mutations are labeled for comparison. **B)** Validation of NFAT-activity screening hits using electroporation of *in vitro* transcribed sgRNAs coupled with ABE8e protein, followed by α-CD3/CD28 stimulation for 16 hours and analysis of GFP (NFAT) or CFP (NFκB) transcriptional activity reporters. * indicate T test p value < 0.05 compared to *HEK*3 relative color control, n=3. **C)** Heatmap of differentially expressed genes between *HEK*3 intergenic mutant control, MAPK1 Y187C, or PHLPP1 S118P edited Jurkat cells activated for 0 or 6 hours as determined by RNA sequencing. The *LCP2* terminating edit is shown for comparison but was not used for statistical testing. **D)** K-means clustering and g:Profiler gene ontology analysis (all p < 0.05) of the differentially expressed gene clusters at 6 hours between *HEK*3 intergenic mutant control, MAPK1 Y187C, or PHLPP1 S118. The *LCP2* terminating edit is shown for comparison but was not used for statistical testing. Cluster numbers count from left to right and are designated by color. **E)** Spinning disk confocal microscopy images showing the subcellular localization of PHLPP1 N-terminal extension (NTE) constructs. “NLS mut” refers to full mutation of the two nuclear localization sequences in PHLPP1’s NTE. The S118P and S118A mutations are also shown. **F)** Quantification of PHLPP1 NTE constructs and the percentage of α-HA signal in the nucleus. * indicates p < 0.05 and ** < 0.01, n=10.

S118 lies within the bipartite nuclear localization sequences (NLSs) at the N-terminus of PHLPP1^63^. To test the hypothesis that phosphorylation controls PHLPP1 subcellular localization, we expressed N-terminal extension (NTE) constructs with the wild type sequence, both halves of the NLSs mutated, or S118P, the result of A-to-G editing. We also included the more archetypal amino acid substitution, S118A, which removes the phosphorylatable residue without potential structural changes introduced by the rotationally constrained amino acid proline. We found that both S118P and S118A reduced PHLPP1 NTE nuclear localization to comparable levels, but less severe than the full NLS mutant (**Figure 5E-F**). These results suggest that phosphorylation of the NTE of PHLPP1 regulates nuclear localization. These data also suggest that serine to proline substitutions are reasonable proxies for loss of a phosphorylatable residue for screening purposes.

### Signaling-to-transcription networks link phosphosites with the expression of specific genes

Transcriptional profiling *HEK3* (intergenic editing control)*, MAPK1* and *PHLPP1* mutant Jurkat T cells showed more differentially expressed genes six hours post stimulus than in resting conditions (BH corrected p value <0.05) (**Figure 5C**), suggesting the mutated phosphosites indeed affect T cell activation-induced transcriptional responses. Inspection of the differentially expressed genes at six hours post T cell activation showed specific T cell-related genes are expressed at subtle but different levels between phosphosite-mutant genotypes (**Figure 6A and Supp Fig 6A**). For example, *BCL11A* and *THEMIS* were activated to a higher extent in MAPK1 Y187C mutants compared to PHLPP1 S118P or control. In contrast, *NR4A3, ZFP36L1,* and *IL21R* were highest in the PHLPP1 S118P cells, whereas *GZMA, TNFSF14, NFKBIA,* and *JUND* were highest in the editing control cells (intergenic *HEK3*). Intracellular GZMB staining 24 hours after T cell activation corroborated the gene expression results, showing a loss of GZMB protein in MAPK1 Y187C cells (**Figure 6B**). To assess the clinical relevance of these genes, we examined the DICE database, which identifies immune cell type-specific expression quantitative trait loci (eQTLs) that double as genome wide association study (GWAS) hits associated with various diseases or human traits^65^. This provides insight into if, and in which immune cells, gene expression modulation could affect T cell-mediated diseases. We found that expression of *CD83*, *BACH2*, *NFKB1*, and *FYN*, which have gene expression dosage effects depending on the phosphosite mutation, increased upon activation of primary human CD4 and CD8 T cells (**Figure 6C**). Moreover, these four genes had multiple GWAS-eQTLs associated with autoimmune diseases (**Figure 6D**). *BACH2* GWAS-eQTLs were specific to T cells and *CD83* was largely attributed to T cells, with some B cell responses (**Supp Fig. 6**). *FYN* was specific to monocytes, whereas *NFKB1* GWAS eQTLs were present in nearly every immune cell tested. Together, these results suggest that mutating different phosphosites in ostensibly the same signaling pathway can alter transcriptional responses, and may elucidate novel points of therapeutic intervention.

**Figure 6.**
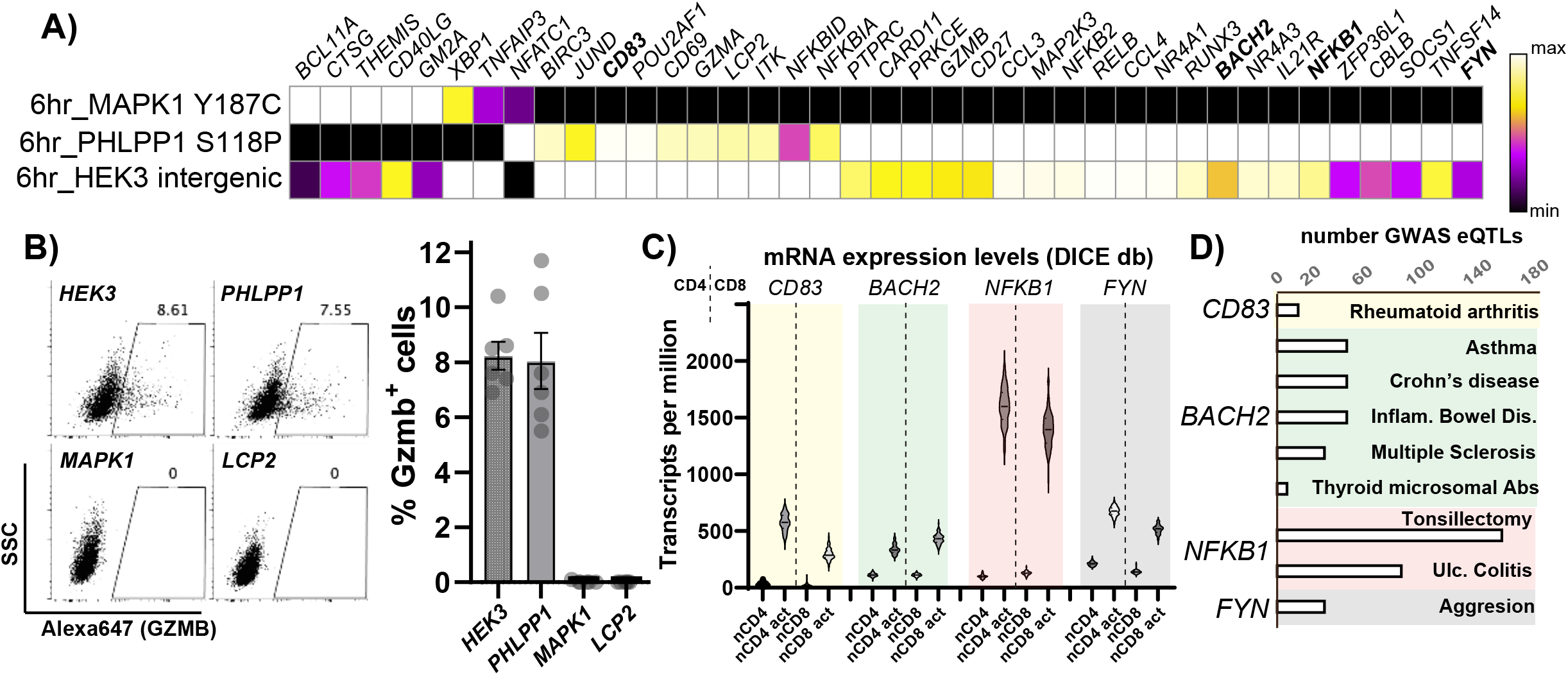
Signaling-to-transcription network mapping links phosphosites with specific genes whose expression levels correlate with human traits. **A)** Select T cell-related differentially expressed genes at six hours post T cell activation between *HEK3* (control), PHLPP1 S118P, and MAPK1 Y187C mutant cells. Bold genes have eQTLs with GWAS hits. **B)** Intracellular GZMB staining in *HEK3* (control), *PHLPP1* S118P, *MAPK1* Y187C or *LCP2* term edited mutant cells 24 hours post-T cell activation. Representative plot (left) and quantification of % GZMB^+^ cells, n = 6. **C)** Violin plot of gene expression in unstimulated or activated (act) naive (n) primary human CD4 or CD8 T cells from the DICE database^65^. **D)** Number of GWAS-eQTLs, and the associated trait or disease, for the four genes that are differentially regulated by different phosphomutants in 6A). “Inflam. Bowel Dis.” indicates inflammatory bowel disease, “Abs” denotes antibodies, and “Ulc. Colitis” indicates ulcerative colitis.

## Discussion

Linking specific signaling events to their downstream functions is a fundamental goal of biology. Choosing novel phosphosites for mechanistic followup studies is often resource intensive and laborious. We aimed to create a screening platform to functionally assess and prioritize individual phosphosites and how they contribute to specific phenotypes in high-throughput by integrating mass spectrometry-based phosphoproteomics with CRISPR-mediated base editor screens. We found that mutation of phosphosites via base editing in a pooled format can assess multiple functional readouts, in positive and negative directions, for proliferative or transcriptional phenotypes. This novel experimental and computational framework will greatly enable future studies, allowing the community to address the complexity of signaling pathways.

Our study focused on phosphosites empirically identified in a parallel experiment rather than from PTM repositories or databases, mitigating the introduction of indiscriminate coding mutations that can have PTM-independent effects^20,21,37^. Moreover, as phosphoproteomic analyses of novel, unstudied systems continue to grow (i.e. patient-specific cancers, primary mouse or human immune cells, etc.) the bioinformatic tools needed to identify base editor-targetable phosphosites from empirical mass spectrometry data will become increasingly important (**Figure 1D**).

We focused on ABE8e for our technology development because it can be readily expressed and purified from *Escherichia coli*, and its activity after electroporation into cells harboring sgRNA-expressing plasmids is highly efficacious (**Figure 2C-D**)^29,30^. Also, as an A-to-G editor it can mutate tyrosine residues, a small but important fraction of total phosphosites. This property is missing from C-to-T editors. The amino acid substitutions made by current base editors to study phosphorylation are not archetypal (S/T to A; Y to F), threonines targeted by ABE8e being the exception (Thr to Ala). Over half of the serines mutated in our study were to proline, a limitation of current base editors. However, since several kinase families are proline directed and the majority of phosphosites are in flexible loops, this is likely to be problematic in only a subset of cases^66–68^. Empirically, the vast majority of Pro substitutions in our data had no effect, arguing it is not inherently disruptive. In fact, our mutational analysis of the PHLPP1 phosphosite showed similar effects between the S118P and S118A mutation, suggesting proline mutations are not invariably detrimental to screen for phosphosite function. Future work will expand testing C-to-T editors, as well as Cas9 mutants that have less restrictive PAM sequences (NG rather than NGG), to expand our targetable phosphosites by 30% and up to 3-fold, respectively.

All gene-centric pathway analyses from genes with mutated phosphosites that altered NFAT activity revealed that the TCR pathway was enriched, indicating the results from our phosphosite base editor screen can recapitulate known aspects of well-studied pathways. Interestingly, gene-level GSEA analysis found “TCR *Calcium* Pathway” enriched in the GFP high bin. The phosphosite mutations driving this result were all in NFAT isoforms, or molecules known to regulate NFAT dephosphorylation: NFATC1/2/3, CABIN1, and RCAN1. This pathway regulates gene expression, cytokine production, and T cell activation through the NFAT signaling pathway by dephosphorylation of NFAT molecules, replicating the effect of Calcineurin (*PPP3CC*) in the translocation of NFAT and T cell activation^48–51^. Site-centric pathway analyses of enriched phosphosite mutations revealed two major trends in the signatures identified in our Jurkat/T cell activation model: TCR signaling and cell cycle. As Jurkat cells are rapidly dividing transformed cells^24^ it is not surprising that many of the phosphosites identified by mass spectrometry, then subsequently in the base editor screens, identified cell cycle genes and phosphorylation events. The DYRK family of kinases is a good example (**Figure 4D**). DYRKs are well known to control NFAT transcriptional activity^69–72^, but also have roles in T cell proliferation^73^. Our results suggest DYRKs have a stronger role in T cell activation compared to proliferation. We envision that applying base editor screens of PTM sites in primary immune cells will provide clearer connections within signaling-to-transcription networks.

Our signaling-to-transcription network mapping identified a novel mechanism for PHLPP1 as a regulator of T cell activation-mediated gene expression. PHLPP1’s function has been primarily studied in cancer contexts^60–62^, though its contribution to regulatory T cell development was identified through genetic studies^64^. Our analyses originally identified it as a positive regulator of NFAT activity, and through transcriptional profiling we found that PHLPP1 can negatively regulate genes downstream of NFκB. This regulation is likely due to the phosphorylation-induced nuclear translocation of PHLPP1, though the precise substrates of dephosphorylation via PHLPP1 remain to be determined. It is worth noting that the phosphopeptide identifying PHLPP1 pS118 was not significantly temporally regulated in our phosphoproteomics data (**Supp Table 1**). This underscores the need for functional screens of PTM sites.

Gene expression profiling of various phosphosite-mutant Jurkats after T cell activation revealed subtle but different transcriptional responses. For example, the *MAPK1* phosphomutant showed decreased expression of *GZMB* compared to the *PHLPP1* mutant, at the protein and mRNA level (**Figure 6**), despite having similar effects on NFAT reporter activity (**Figure 5**). *MAPK1* phosphomutant cells expressed less *NR4A1* and *NR4A3* than the PHLPP1 phosphomutant cells. GZMB is the major effector molecule of the cytotoxic program^74^ and NR4As have been shown to be drivers of T cell exhaustion^75,76^. We also found that through modifying signaling through MAPK1 and PHLPP1, expression levels of genes associated with autoimmunity are altered (**Fig. 6D**). These results together lead to an intriguing possibility of fine-tuning expression levels of specific genes, through genetic or small molecule inhibitor manipulation of signaling pathways, to improve adoptive cell transfer and cancer immunotherapies, or to modulate autoimmune disorders.

Proteome-wide, PTM-centric base editing coupled to phenotypic screens provides a powerful experimental framework to untangle the vast network of biochemical signaling reactions and how they lead to the control of specific cellular functions. This approach directly assesses the impact of a phosphosite mutation on a given phenotype rather than relying purely on evolutionary or structural conservation, and provides interpretable results with high confidence for further mechanistic studies^37,77–80^. With currently available base editors it should be possible to functionally screen other PTMs including acetylation/ubiquitination/methylation (lysine)^81^, O-GlcNAcylation (Ser/Thr)^82^, cysteine^83^, or even specific proteolysis events (caspases)^84^, utilizing established infrastructure common to many research institutions. We envision this approach to be widely enabling to the cell biology community.

## Methods

### Cell Culture

Jurkat E6.1 and HEK293T cells were purchased through ATCC. Triple parameter reporter (TPR) Jurkat cells^36^ were purchased from Professor Peter Steinberger at the Medical University of Vienna. TPR Jurkat cells and E6.1 Jurkat cells were cultured and passaged in Roswell Park Memorial Institute medium 1640 (RPMI-1640) plus GlutaMax (Thermo Fisher Scientific) supplemented with 10% heat inactivated fetal bovine serum. Cells were passaged and maintained at cell densities between 1-5e5 cells per mL.

HEK293T cells were cultured and passaged in Dulbecco’s Modified Eagle Medium (DMEM) supplemented with 10% heat inactivated fetal bovine serum.

All cells were incubated and maintained at 37°C with 5% CO2.

### Phosphoproteomic analysis

Jurkat E6.1 cells (ATCC) were activated in 96 well tissue culture plates for the stated times at 2e5 cells/mL. Plates were coated with 3.33 μg/mL α-CD3 (HITa3) and α-CD28 antibodies (Biolegend), in 100 μL PBS at 4°C overnight, followed by one cold PBS wash. To stop the reaction, the cells were transferred to ice cold PBS, washed twice, flash frozen, and stored at -80°C until processing. Phosphoproteomic analysis was performed as previously described^25,26^. Data was analyzed using Spectrum Mill (Agilent and Broad Institute) for phosphopeptide identification and quantification. The TMT denominator for each sample was the mean of all TMT channels. After global median normalization and median absolute deviation scaling, a moderated F test (limma, R) was performed to identify “regulated” phosphopeptide levels across the time series. Statistically significant phosphopeptides (Benjamini-Hochberg adj. p value <0.05) were visualized using Morpheus (https://software.broadinstitute.org/morpheus/). The .json file associated with this manuscript, **Supplemental Data File 1**, can be used to explore these data in Morpheus.

### ABE8e protein expression and purification

Recombinant ABE8e was expressed and purified as previously described^30^. Briefly, ABE8e was expressed as an 8xHis tagged protein from a rhamnose-inducible promoter in BL21-Star DE3 cells with low RNase activity (Thermo Fisher Scientific). At an OD600 of ∼0.8, the terrific broth *Escherichia coli* cultures were cold shocked on ice for one hour, and induced with 0.8% final concentration of rhamnose. Roughly 24 hours later, cells were lysed via lysozyme and sonication, and ABE8e protein was purified on Ni-NTA resin. After imidazole elution, ABE8e protein was further purified using cation exchange. Fractions were monitored by UV and SDS-PAGE. ABE8e protein containing fractions were pooled, concentrated to ∼90 pmol/μL using 100 MW cutoff filters, aliquoted, flash frozen, and stored at -80°C.

### Base editing with arrayed lentivirus

Oligonucleotides containing the protospacer were chosen by hand^85^ or from our original bioinformatic analysis, and ordered from IDT DNA technologies. CCACG was added to the forward oligo, where AAAC was added to the reverse complement of the protospacer sequence. An A on the 3’ end of the reverse complement oligo was also appended. pRDA118 (Addgene Plasmid # 133459) was digested with BsbmI_V2 (NEB) and FastAP (Thermo) for the last five minutes, followed by gel purification. Oligos were mixed 1:1 at 100 μM, phosphorylated via PNK (NEB) and annealed after a 5 minute 95°C step at 5°C per five minutes until 25°C. Annealed oligos were diluted 1:200 and 1 μL was mixed with 25-50 ng of digested backbone. T4 ligation (NEB) was performed for 20 minutes at 37°C and the 5-10 μL ligation reaction was transformed into Stbl3 *E. coli*, made in house. Colonies were verified via Sanger sequencing. One μg of sgRNA plasmids were transfected with Lipofectamine 3000 (Invitrogen) 1 μg of pPAX2 and 0.1 μg VSVG into HEK293Ts seeded the night before at 2e5 cells per 6 well. After three hours the 2.5 mL media was replaced with 5 mLs media supplemented with 1% BSA. Supernatants were collected after 3 days, filtered to remove HEK293T cells, concentrated 10-fold with Lenti-X precipitation solution (Alstem), and aliquots were flash frozen and stored at -80°C.

Wild-type or TPR Jurkat cells were spinfected for two hours at 666 x g. Two days later, 2 μg/mL puromycin was added until the non-transduced cells were all dead (2-4 days). To test base editor delivery molecules, only the *HEK3* site-targeting sgRNA was used, and electroporation was performed using Lonza Nucleofection with the SE or P3 nucleofection solution. pCMV-BE4max was used for plasmid DNA. Base editor mRNAs were generated by *in vitro* transcription using the HiScribe T7 High-Yield RNA synthesis kit (NEB Cat No. E2040S) via the method described previously^86^. NEBnext polymerase was used to PCR-amplify template plasmids and install a functional T7 promoter and a 120 nucleotide polyadenine tail. Transcription reactions were set up with complete substitution of uracil by N1-methylpseudouridine (Trilink BioTechnologies Cat No. N-1080) and co-transcriptional 5’ capping with the CleanCap AG analog (Trilink BioTechnologies Cat No. N-7113) to generate a 5’ Cap1 structure. mRNAs were purified using ethanol precipitation according to kit instructions, dissolved in nuclease-free water, and normalized to a concentration of 2 micrograms per microliter using Nanodrop RNA quantification of diluted test samples.

### Base editing with *in vitro* transcribed guides and ABE8e protein

sgRNAs were transcribed *in vitro* using the EnGen sgRNA Synthesis kit (NEB) with oligonucleotides containing the protospacers. The sgRNAs were then purified (NEB Monarch RNA Cleanup kit) and quantified. 3 μg of purified sgRNA were then mixed with 1 μL of 90 μM ABE8e protein in Lonza P3 electroporation buffer in a total reaction volume of 10 μL and incubated at room temperature to allow ribonucleoprotein complex to form. TPR Jurkat cells were collected, spun, washed twice with 37°C PBS, and 2e5 cells were resuspended per cuvette well in Lonza electroporation buffer P3 with the ABE8e and sgRNA ribonucleoprotein complex to a total reaction volume of 20 μL. Electroporation and cell recovery was performed according to the manufacturer’s instructions. Two days post nucleofection, gDNA of the cells was extracted, and base editing efficiency of each population was determined through PCR amplification of the edited genomic region of interest and quantified through Sanger sequencing and EditR software analysis^31^. To produce purely edited cell populations, single cell clones were isolated from the corresponding bulk edited cell populations by diluting 0.8 cells per well in 96-well plates. Once isolated, these cells were grown until confluence, then individually genotyped through extraction of its gDNA and PCR amplification of the edited genomic region of interest. Once validated, 4-8 single cell clones were then mixed together to avoid clone-specific effects.

### Phosphosite library design and construction

All phosphorylation sites determined through the accompanying phosphoproteomic analysis were filtered for phosphosites that were localized in the peptide with a confidence of 90% or greater. For all genes with phosphosites, we used an in-house base editor design tool to first design all possible guides targeting these genes. All adenines in the window of 4-8 of the sgRNA (where 1 is the most PAM-distal position, and positions 21-23 are the PAM) were considered to be edited. sgRNAs with cloning sites, poly Ts and greater than five perfect matches in the genome were excluded. sgRNAs targeting the phosphosite of interest were then picked. sgRNAs predicted to make silent edits at phosphosites but also predicted to have bystander edits in the window of 4-8 were excluded. We then included 250 non-targeting controls, 250 intergenic controls, 250 controls targeting splice sites of essential genes and 250 controls targeting splice sites of TCR genes. The code is available at https://github.com/mhegde/base-editor-design-tool.

### Proteome-wide base editor screen

TPR Jurkat cells were spinfected in 4 µg/mL polybrene at 30°C for 45 minutes at 666 x g at a multiplicity of infection of 0.3, assuming 30% transduction efficiency, and maintained a 500x library coverage. Transduction quadruplicates were used for downstream replicates. Puromycin was added to 2 μg/mL and cells were selected for stable integrants for seven days. Three days after removal of puromycin, an aliquot of cells corresponding to 500x library coverage was saved as the pre-ABE8e inputs. Cells were washed with 37°C PBS twice, and resuspended in SE nucleofection reagent (Lonza) with 94 pmoles of ABE8e protein per well. Each well was 2e5 library-containing TPR cells in 19 µL of SE + 1 µL ABE8e protein, and all 16 wells were used simultaneously for a library coverage of 250x. Electroporation and cell recovery was performed according to the manufacturer’s instructions. Six days after ABE8e protein introduction, another 500x aliquot of cells was frozen for the post-ABE8e sample.

14 days after ABE8e protein introduction, the mutant library TPR Jurkat pool was washed in room temp PBS, and activated as described above. 2000x library coverage of TPR cells was prepared for FACS. FACS was performed on a Bigfoot Spectral Cell Sorter (ThermoFisher) and roughly 1e6 cells were collected per bin. The top one, and bottom two 12.5% bins were sorted. The two bottom bins (“bottom” and “low”) were sorted to avoid unactivated cells, which always was about 15% of all cells. Only the second lowest bin (“low”) was used for downstream analyses due to the superior performance of controls, and to ensure cells were activated.

Collected cells were pelleted and stored at -80°C until further processing. Genomic DNA (gDNA) was isolated and PCR-amplified for barcode abundance determination as previously described^20^. Standard Illumina adapters were added and stagger regions were introduced for base diversity, according to the protocol “sgRNA/shRNA/ORF PCR for Illumina Sequencing” (The Broad Institute GPP). Libraries were sequenced at a depth of 10e6 reads or more at AUGenomics.

### Differential sgRNA abundance analyses via MAGeCK and MAGeCKFlute Programs

To identify the functional phosphorylation sites through mutational analyses, we analyzed raw reads of GFPhigh and GFPlow samples using the MAGeCK program (v0.5.9.5), used for analyzing CRISPR screens^17^. To summarize the results of MAGeCK’s phosphosite and sgRNA data, we used MAGeCKFlute program (version 2.4.0)^46^. Before conducting the pathway enrichment and gene-centric GSEA analysis with MAGeCKFlute, we converted the phosphorylated and mutated gene products to their corresponding gene symbols. For site-centric analyses (PTM-SEA, Kinase Library), we used the MAGeCK output for ssGSEA2.0 R package with default settings^27^. PTM-SEA included iKiP data^87^ but excluded the LINCS P100^88^ terms for clarity. To investigate the putative kinases responsible for phosphorylating phosphosites abundant in post-base editing and NFAT signaling, we employed The Kinase Library Enrichment analysis, which offers an atlas of primary sequence substrate preferences for the human serine/threonine kinome, was used on the PhosphositePlus.org website^45^.

### PHLPP1 localization

Plasmids encoding the N-terminal extension (NTE) of PHLPP1 were HA tagged^63^, and the appropriate codons were mutated using site-directed mutagenesis (Agilent). HeLa cells were transiently transfected with WT, NLS mutant, S118P, or S118A PHLPP1 NTEs, and after two days were fixed, and stained with anti-HA antibody (Cell Signaling, 3742), Alexa Fluor-647-phalloidin (Invitrogen, A22285), and DAPI stains. Images were acquired using spinning disk confocal microscopy. Quantification was performed using individual cells and were statistically tested using a one-way ANOVA with Tukey’s multiple test corrections.

### Transcriptional profiling of phosphosite-mutant T cell lines

The various phosphosite mutants were introduced into TPR Jurkat cells via *in vitro* transcribed sgRNAs described above. After single cell cloning 4-8 single cell clones were mixed together to avoid clone-specific effects. Cells were incubated and activated with α-CD3/CD28 agonist antibodies as described above, except that the antibody concentrations were 1 μg/mL. Cells were activated for 0 and six hours.

Cells were stained with BioLegend TotalSeq-C Human Universal Cocktail and anti-human hashtag antibodies according to NYCG CITEseq protocols [http://cite-seq.com/] and processed using 10X Genomics 5’ HT with Feature Barcode assay according to protocol, with the addition of selective transcript removal using Jumpcode Genomics CRISPRclean Single Cell Boost kit.

Briefly, 100K cells from each phosphosite-mutant line per time point were blocked with BioLegend Human TruStain FcX Fc receptor blocking solution. Cells were then stained with BioLegend TotalSeq-C Human Universal Cocktail resuspended in BioLegend Cell Staining Buffer (CSB) at a concentration of 1 vial per 500K cells and anti-human hashtag antibodies at a concentration of 0.75 μg/1e6 cells for 30 minutes at 4°C. Cells were then washed 3x in CSB. Cell viability and concentration was assessed using the Moxi Go II and Moxi Cyte Viability Reagent containing propidium iodide (PI). Cells were pelleted and resuspended in 0.04% BSA in PBS for a final concentration of 1300-1600 cells/μl and processed using 10X Genomics Next GEM Single Cell 5’ HT v2 assay. Gene Expression (GEX) and Cell Surface Protein (CSP) libraries were constructed according to protocol (CG000424 Rev D) with the following deviation: Post-ligation product of the GEX library was subjected to Jumpcode Genomics CRISPRclean Single Cell Boost kit following protocol. Libraries were sequenced at a targeted depth of 50K reads/cell for GEX and 5k reads/cell for CSP on an Illumina NovaSeq 6000 and an Element Aviti. Cellranger count v7.1.0 was used to generate cloupe files for analysis.

Data was analyzed using the Loupe Browser (10X Genomics). For Figure 5C, *HEK3*, MAPK1, *LCP2*, and *PHLPP1* phosphosite-mutant cells were selected for zero and six hours post-activation, and differential gene expression (log2 fold change) was determined using local (sample specific) expression. Genes with a p value less or equal to 0.1 were determined to be regulated, as recommended by the software. For Figure 5D, differentially expressed genes were determined using only *HEK3*, MAPK1, and *PHLPP1* cells, though *LCP2* terminating edit cells’ log2 fold change values were included in the plot for context. For Figure 6A, genes were selected from the plot in 5D through their known involvement in T cell signaling.

### Granzyme B staining

3.5e5 cells were incubated with fixable viability dye prior to fixation and permeabilization using Ghost Dye UV 450 (Cytek) at a 1:100 dilution, in a total volume of 50μL, for 15 minutes in PBS containing 2% FBS (PBS 2%) at 4°C and then washed once in the same medium at 500g for 4 minutes. Cells were fixed in 50 μL of BD Cytofix during 20 minutes at 4°C and then washed in PBS 2% at 500g for 4 minutes. After, cells were fixed and permeabilized using the eBioscience Foxp3 / Transcription Factor Staining Buffer Set (Thermo Fisher) for 40 minutes at 4°C and then washed in perm/wash buffer at 500g for 4 minutes. Expression of granzyme b was evaluated using the Granzyme B-A647 monoclonal antibody (clone GB11, BD biosciences) at a 1:100 dilution, in a total volume of 50μL. Cells were incubated 1 hour at RT followed by a 4°C incubation overnight. Cells were washed in perm/wash at 500g for 4 minutes and then resuspended for analysis. Analyses were performed on LSRII cytometer (BD Biosciences). 10,000 events were recorded and data analyses were performed in FlowJo software (Tree Star, Ashland, OR).

## Acknowledgements

We thank Adam Haber, Jeff Johnson, Pandurangan Vijayanand, Egle Kvedaraite, B. Hamilton, Adam Rubin, Tomer M. Yaron, and Matteo Gentili for useful discussion. We also thank Gary and Shuko Clouse for support, Dr. Peng Guo and the Nikon Imaging Center at UCSD for the support on microscopy experiments, and Steven Carr, Senior Director of Proteomics Broad Institute of MIT and Harvard for preliminary discussions, guidance, support, and resources. This work was supported by NIH NIGMS R35GM147554 (SAM), NIH R35 GM122523 (ACN), NIH U01AI142756, R35GM118062, RM1HG009490 and HHMI (DRL), Stem Cell Network Jump Start Award (ECR-C4R1-7) for CGD who is a Michael Smith Health Research BC Scholar. ACJ was supported in part by the UCSD Graduate Training Program in Cellular and Molecular Pharmacology (T32 GM007752) and National Science Foundation Graduate Research Fellowship Program (DGE-1650112). PGH is supported by NIH grants AI 040127 and AI 109842, The NovaSeq 6000 was acquired through the Shared Instrumentation Grant (SIG) Program (S10) S10OD025052; La Jolla Institute for Immunology Next Generation Sequencing Core Facility RRID:SCR_023107. FACSAria-3 was acquired through the Shared Instrumentation Grant (SIG) Program (S10): RR027366; La Jolla Institute for Immunology Flow Cytometry Core RRID:SCR_014832. This article is subject to HHMI’s Open Access to Publications policy. HHMI lab heads have previously granted a nonexclusive CC BY 4.0 license to the public and a sublicensable license to HHMI in their research articles. Pursuant to those licenses, the author-accepted manuscript of this article can be made freely available under a CC BY 4.0 license immediately upon publication.

DRL is a consultant and/or equity owner for Prime Medicine, Beam Therapeutics, Pairwise Plants, Chroma Medicine, and Nvelop Therapeutics, companies that use or deliver genome editing or epigenome engineering agents. The remaining authors have no conflicts of interest.

## Data Availability

Raw mass spectrometry data and metadata can be accessed at ftp://MSV000092965@massive.ucsd.edu. Raw RNA sequencing data can be accessed at GEO accession ID GSE244164.

## Code Availability

The code for the base editor design tool is available at https://github.com/mhegde/base-editor-design-tool.

## Author contributions (CRediT)

Conceptualization: SAM. Methodology: PHK, ACJ, MH, CdB, GN, SAM. Software: MH, JGD. Validation: PHK, AADS, AJC, SAM. Formal Analysis: AADS, MEO, RB, SA, RAG, GN, SAM. Investigation: PHK, AADS, MB, ACJ, MEO, MIM, NP, PGH, RB, ACN, SA, RAG, CdB, GN, SAM. Resources: MB, NP, JL, PGH, DRL, JGD, GN, CdB, and SAM. Data Curation: AADS, MEO, MH, MIM, RB, JGD, SA, RAG, and SAM. Writing: All authors. Visualization: PHK, AADS, AJC, MEO, SA, and SAM. Supervision: SAM. Project Administration: SAM. Funding Acquisition: DRL, SAC, and SAM.

## Figures legends

**Supplemental Figure associated with Figure 3.**
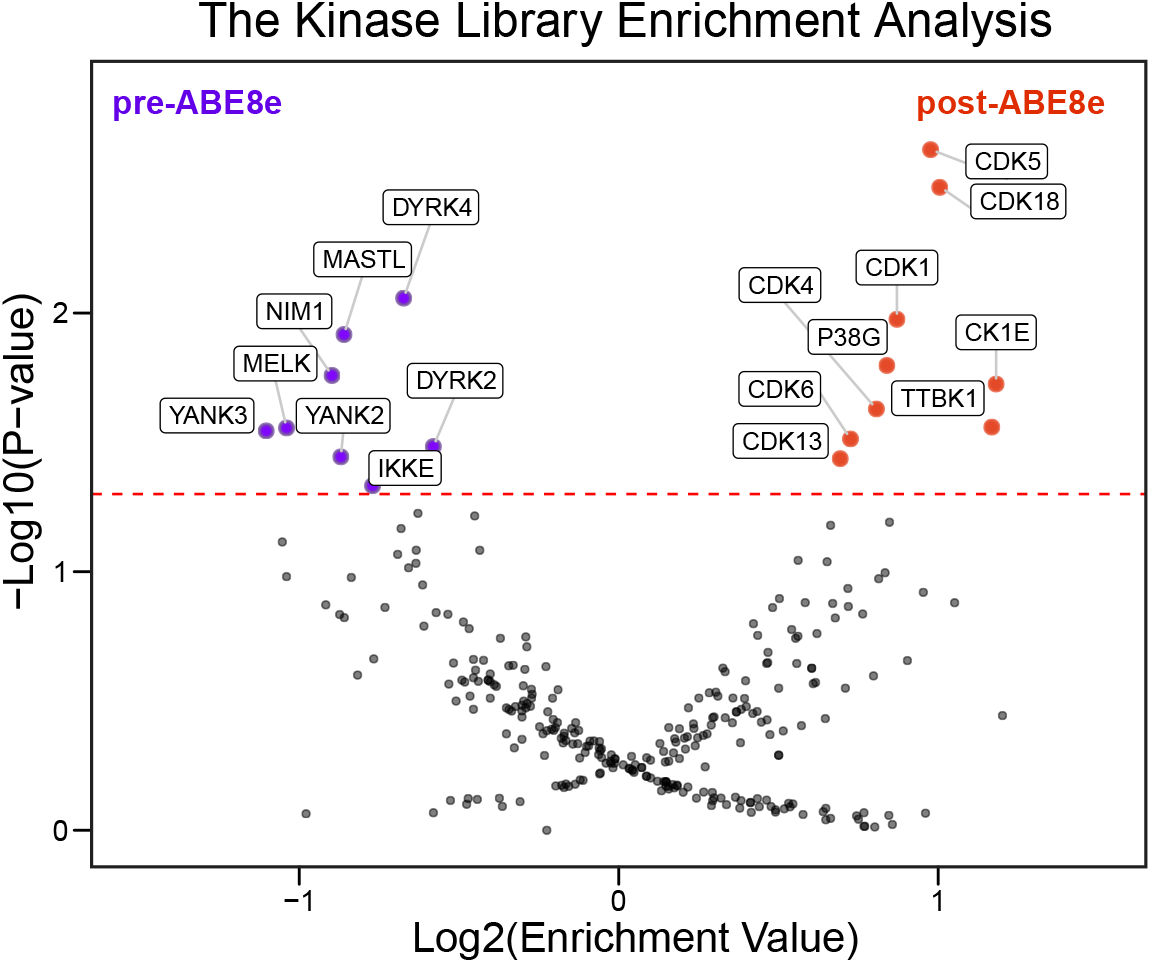
Kinase Library, site-centric enrichment analysis of phosphosite mutants, as an aggregate motif, enriched in post ABE8e-edited cells (purple) or pre ABE8e-edited cells (red) bins. Enrichment Values were determined by MAGeCK analysis.

**Supplemental Figure associated with Figure 4.**
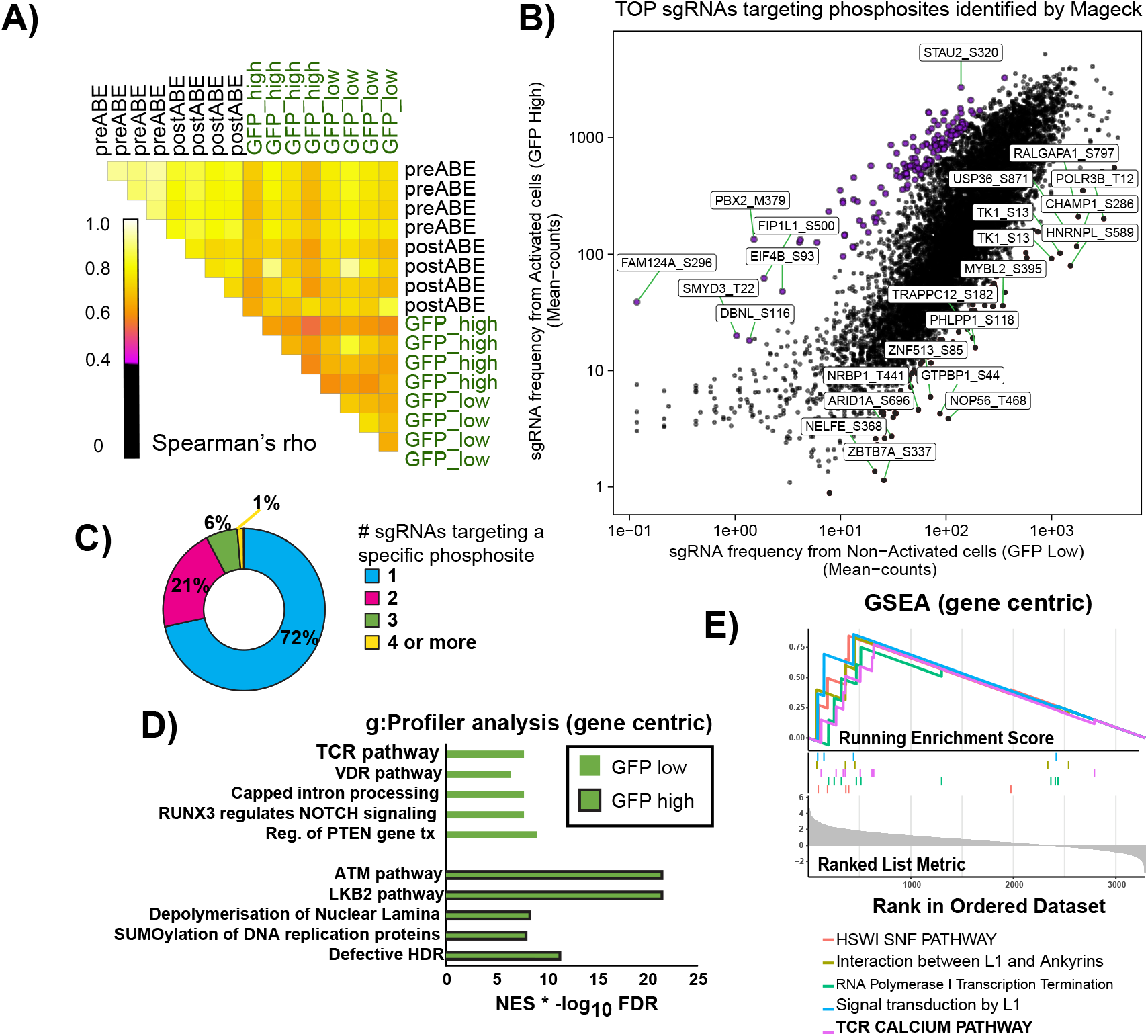
Quality control and characterization of phosphosite base editing coupled to NFAT activity reporters. **A)** Pairwise Spearman correlations between all normalized log transformed read counts across replicates and experimental conditions. 0.4 is the lower limit cut off in black. **B)** Mean (across replicates) sgRNA counts for individual sgRNAs prior to collapsing redundant phosphosite targets. **C)** Percentage of phosphosite targets with one or more protospacer sequences. **D)** g:Profiler analysis (gene-centric) of genes with phosphosite mutations enriched in the GFP low or GFP high bins. For the x-axis the normalized enrichment score (NES) was multiplied by the -log10 FDR **E**) GSEA (gene-centric) analysis of gene sets enriched in the GFP high bin. TCR Calcium Pathway is bolded.

**Supplemental Figure associated with Figure 6.**
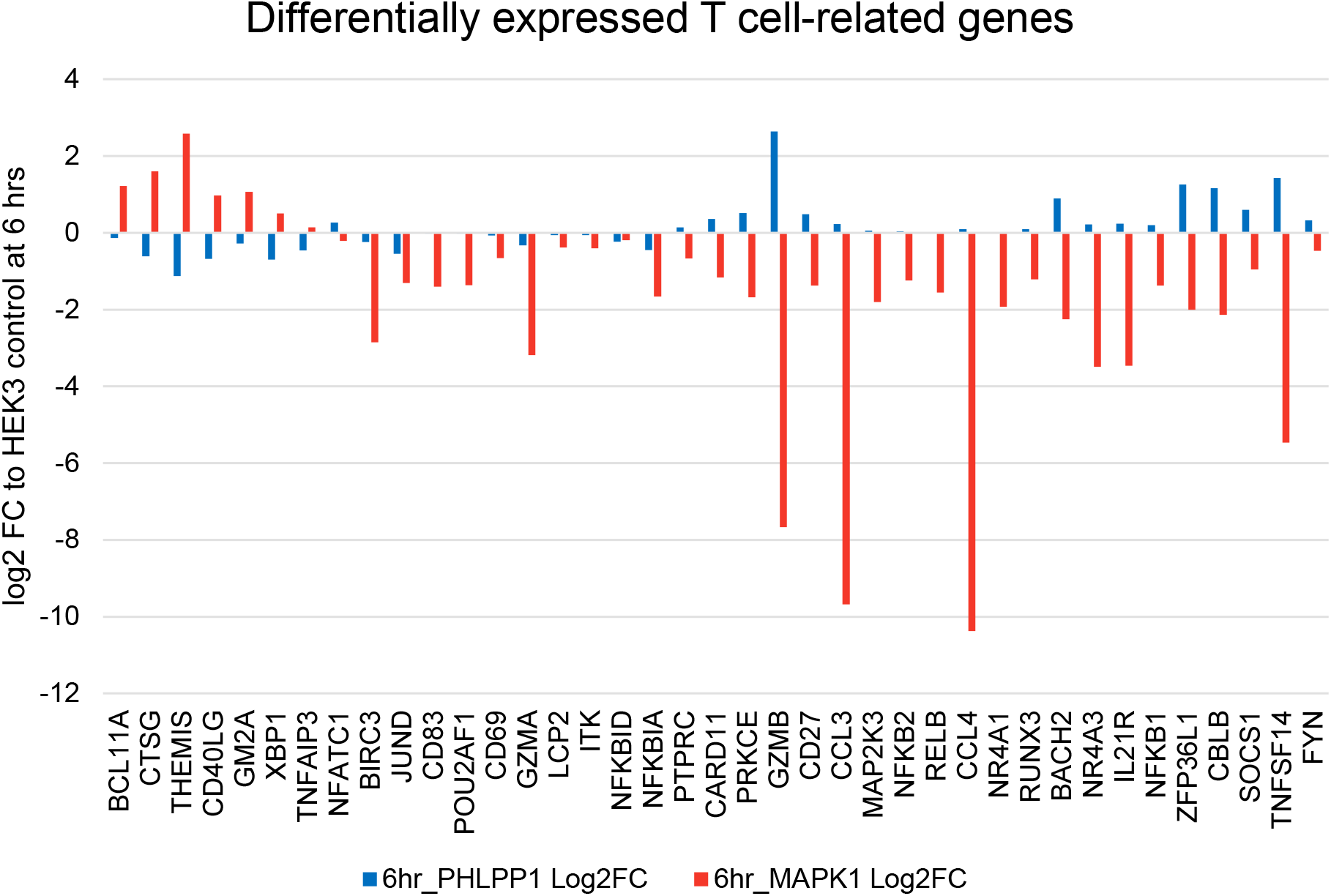
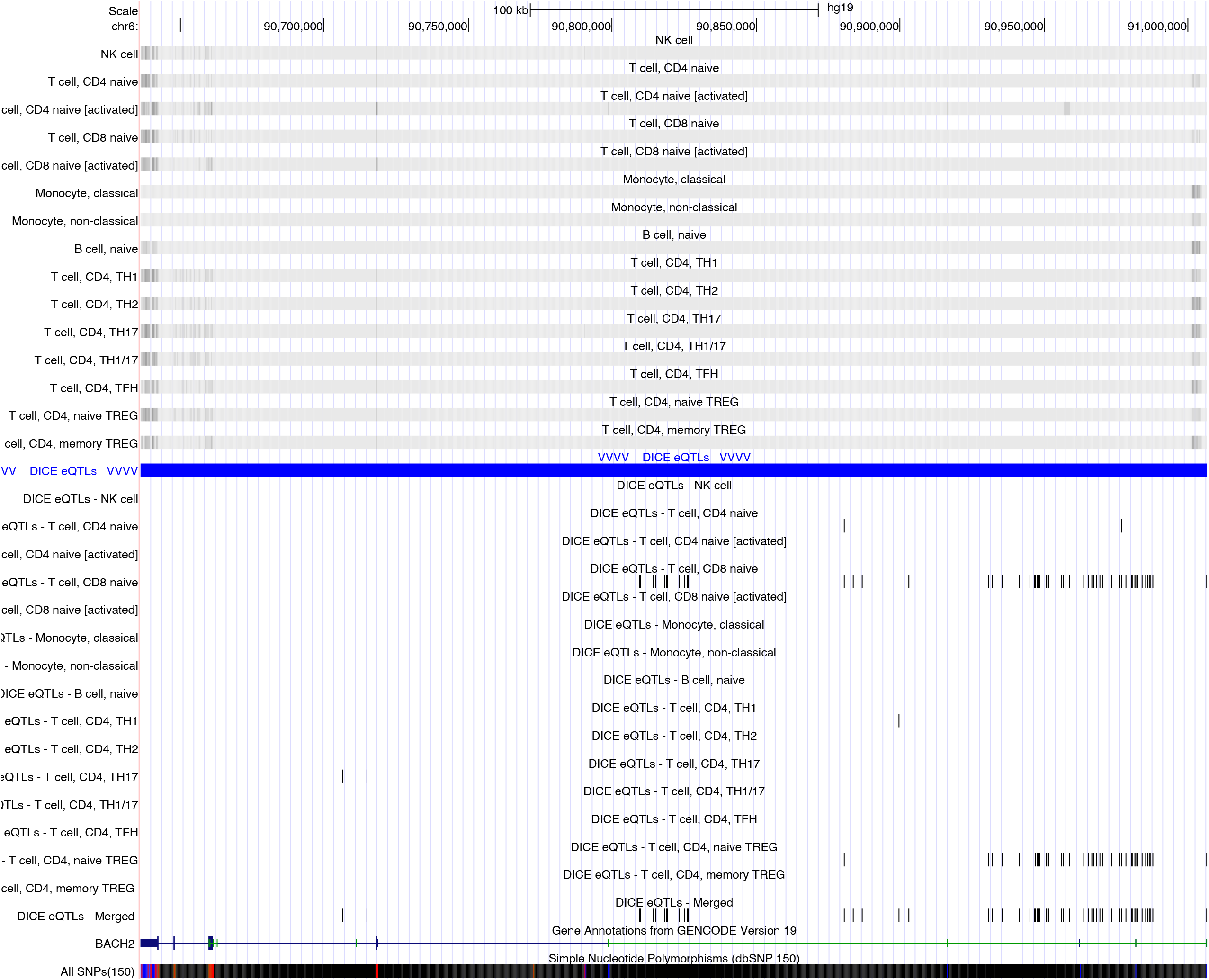
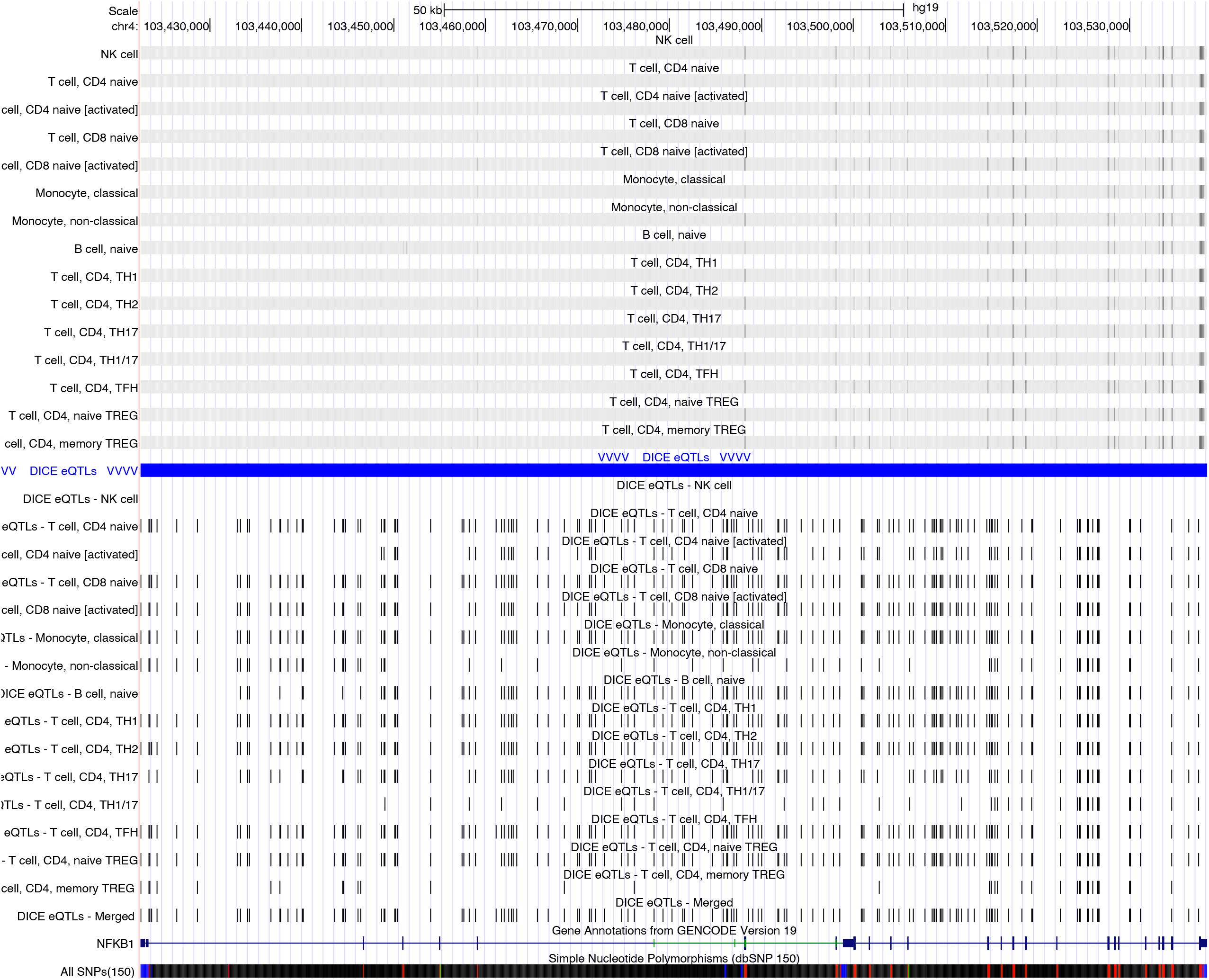
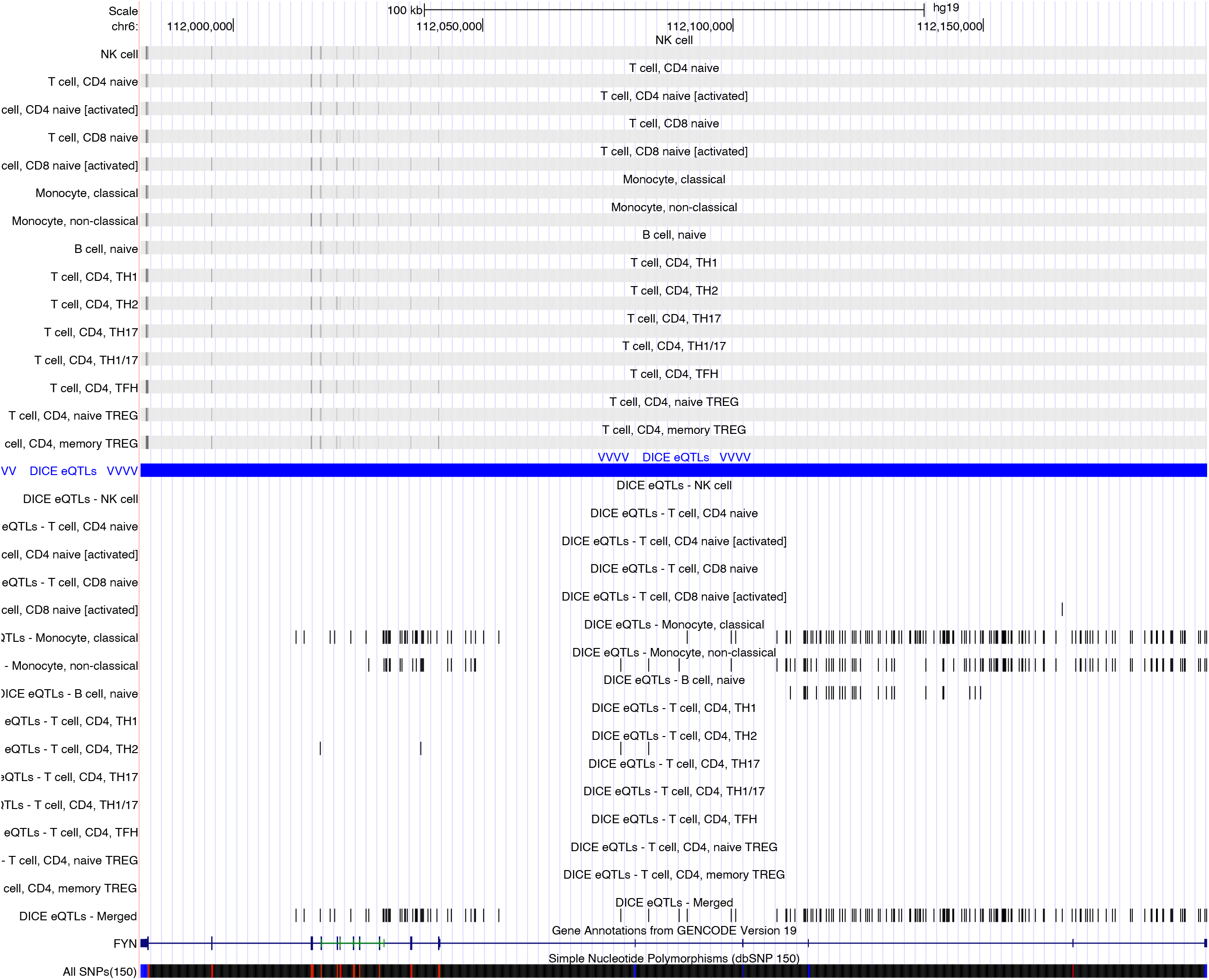
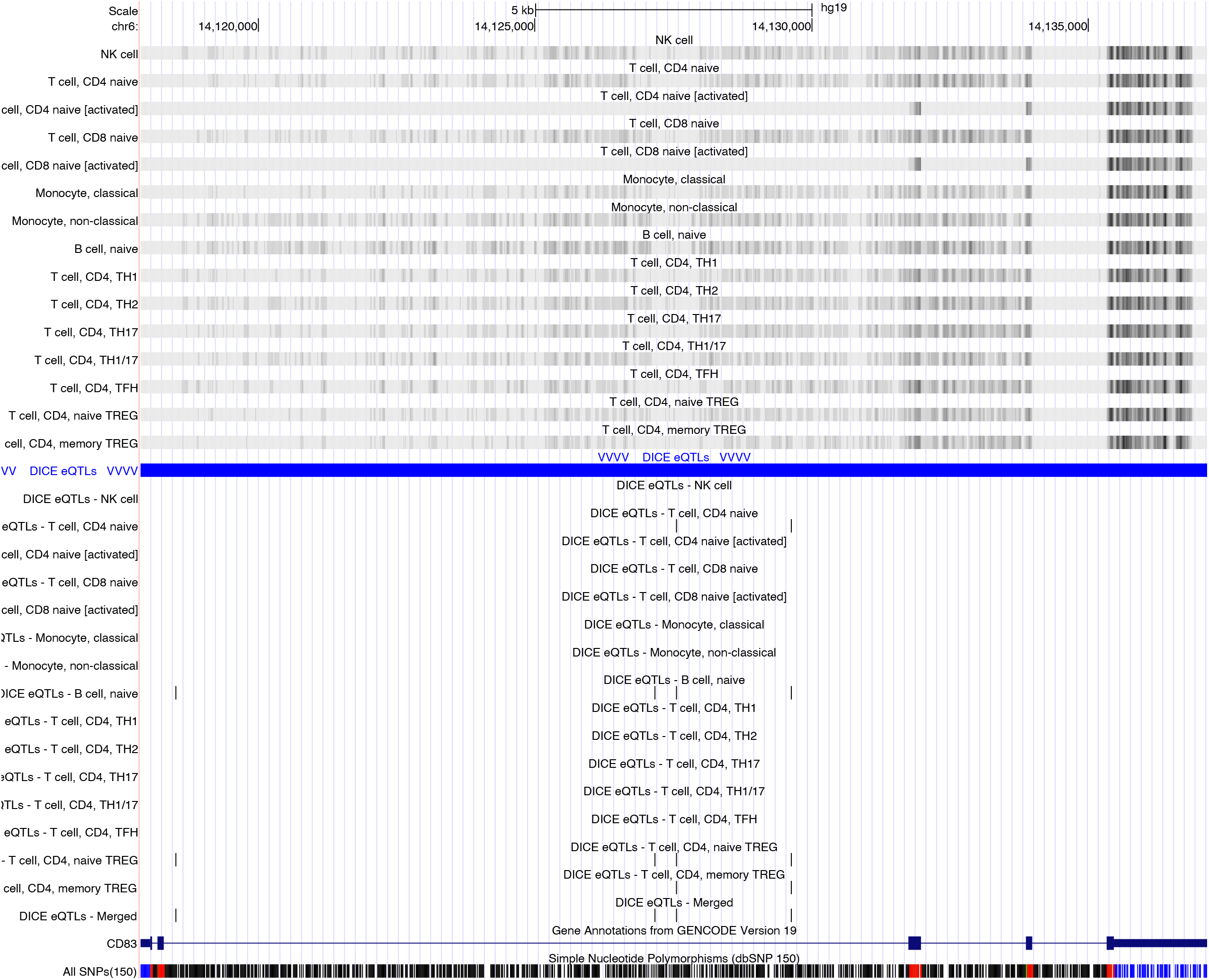
**A)** Log2 fold change of select T cell genes differentially expressed between PHLPP1 S118P and MAPK1 Y187C mutant cells, compared to *HEK3* control cells. **B)** UCSC genome browser tracks derived from the DICE database showing eQTLs associated with specific genes (bottom left) across various cell types (under blue bar). *BACH2, NFKB1, FYN*, and *CD83* are shown in that order.

## Supplemental Information

**Supplemental Table 1.** Phosphoproteomics analyses processed by Spectrum Mill and statistically tested by Protigy. “modF” provides all analysis results and measurement values. “Class vector” provides sample key for TMT channels. “Description of table header” refers to modF and describes where which analysis comes from.

**Supplemental Table 2A.** ABE8e-targetable phosphosites.

**Supplemental Table 2B.** BE4-targetable phosphosites

**Supplemental Table 3.** Differential analysis of sgRNA abundances between pre- and post-ABE8e protein introduction

**Supplemental Table 4.** Differential analysis of sgRNA abundances between GFP high and GFP low bins--NFAT activity screen

**Supplemental Table 5.** RNA-seq analysis of activated Jurkat T cells with various phosphosite mutations.

## Notes

ftp://

## References

1 Hunter T. Why nature chose phosphate to modify proteins. Philos Trans R Soc Lond B Biol Sci 2012;367:2513–6. 10.1098/rstb.2012.0013.

2 Manning G, Whyte DB, Martinez R, Hunter T, Sudarsanam S. The protein kinase complement of the human genome. Science 2002;298:1912–34. 10.1126/science.1075762.

3 Chen MJ, Dixon JE, Manning G. Genomics and evolution of protein phosphatases. Sci Signal 2017;10.: 10.1126/scisignal.aag1796.

4 Krug K, Jaehnig EJ, Satpathy S, Blumenberg L, Karpova A, Anurag M, et al. Proteogenomic Landscape of Breast Cancer Tumorigenesis and Targeted Therapy. Cell 2020;183:1436–56.e31. 10.1016/j.cell.2020.10.036.

5 Katrancha SM, Shaw JE, Zhao AY, Myers SA, Cocco AR, Jeng AT, et al. Trio Haploinsufficiency Causes Neurodevelopmental Disease-Associated Deficits. Cell Rep 2019;26:2805–17.e9. 10.1016/j.celrep.2019.02.022.

6 Martinez-Val A, Bekker-Jensen DB, Steigerwald S, Koenig C, Østergaard O, Mehta A, et al. Spatial-proteomics reveals phospho-signaling dynamics at subcellular resolution. Nat Commun 2021;12:7113. 10.1038/s41467-021-27398-y.

7 Bouhaddou M, Memon D, Meyer B, White KM, Rezelj VV, Correa Marrero M, et al. The Global Phosphorylation Landscape of SARS-CoV-2 Infection. Cell 2020;182:685–712.e19. 10.1016/j.cell.2020.06.034.

8 Koch H, Wilhelm M, Ruprecht B, Beck S, Frejno M, Klaeger S, et al. Phosphoproteome profiling reveals molecular mechanisms of growth-factor-mediated kinase inhibitor resistance in EGFR-overexpressing cancer cells. J Proteome Res 2016;15:4490–504. 10.1021/acs.jproteome.6b00621.

9 Zecha J, Gabriel W, Spallek R, Chang Y-C, Mergner J, Wilhelm M, et al. Linking post-translational modifications and protein turnover by site-resolved protein turnover profiling. Nat Commun 2022;13:165. 10.1038/s41467-021-27639-0.

10 Paulo JA, Gygi SP. A comprehensive proteomic and phosphoproteomic analysis of yeast deletion mutants of 14-3-3 orthologs and associated effects of rapamycin. Proteomics 2015;15:474–86. 10.1002/pmic.201400155.

11 Needham EJ, Parker BL, Burykin T, James DE, Humphrey SJ. Illuminating the dark phosphoproteome. Sci Signal 2019;12:eaau8645. 10.1126/scisignal.aau8645.

12 Hornbeck PV, Zhang B, Murray B, Kornhauser JM, Latham V, Skrzypek E. PhosphoSitePlus, 2014: mutations, PTMs and recalibrations. Nucleic Acids Res 2015;43:D512–20. 10.1093/nar/gku1267.

13 Dixit A, Parnas O, Li B, Chen J, Fulco CP, Jerby-Arnon L, et al. Perturb-Seq: Dissecting Molecular Circuits with Scalable Single-Cell RNA Profiling of Pooled Genetic Screens. Cell 2016;167:1853–66.e17. 10.1016/j.cell.2016.11.038.

14 Parnas O, Jovanovic M, Eisenhaure TM, Herbst RH, Dixit A, Ye CJ, et al. A Genome-wide CRISPR Screen in Primary Immune Cells to Dissect Regulatory Networks. Cell 2015;162:675–86. 10.1016/j.cell.2015.06.059.

15 Shifrut E, Carnevale J, Tobin V, Roth TL, Woo JM, Bui CT, et al. Genome-wide CRISPR screens in primary human T cells reveal key regulators of immune function. Cell 2018;175:1958–71.e15. 10.1016/j.cell.2018.10.024.

16 Shalem O, Sanjana NE, Zhang F. High-throughput functional genomics using CRISPR–Cas9. Nat Rev Genet 2015;16:299–311. 10.1038/nrg3899.

17 Li W, Xu H, Xiao T, Cong L, Love MI, Zhang F, et al. MAGeCK enables robust identification of essential genes from genome-scale CRISPR/Cas9 knockout screens. Genome Biol 2014;15:554. 10.1186/s13059-014-0554-4.

18 Meyers RM, Bryan JG, McFarland JM, Weir BA, Sizemore AE, Xu H, et al. Computational correction of copy number effect improves specificity of CRISPR–Cas9 essentiality screens in cancer cells. Nat Genet 2017;49:1779–84. 10.1038/ng.3984.

19 Rees HA, Liu DR. Base editing: precision chemistry on the genome and transcriptome of living cells. Nat Rev Genet 2018;19:770–88. 10.1038/s41576-018-0059-1.

20. Hanna RE, Hegde M, Fagre CR, DeWeirdt PC, Sangree AK, Szegletes Z, et al. Massively parallel assessment of human variants with base editor screens. Cell 2021;184:1064–80.e20. 10.1016/j.cell.2021.01.012.

21. Lue NZ, Garcia EM, Ngan KC, Lee C, Doench JG, Liau BB. Base editor scanning charts the DNMT3A activity landscape. Nat Chem Biol 2022. 10.1038/s41589-022-01167-4.

22. Xu P, Liu Z, Liu Y, Ma H, Xu Y, Bao Y, et al. Genome-wide interrogation of gene functions through base editor screens empowered by barcoded sgRNAs n.d. 10.21203/rs.3.rs-57831/v1.

23 Yeh W-H, Chiang H, Rees HA, Edge ASB, Liu DR. In vivo base editing of post-mitotic sensory cells. Nat Commun 2018;9:2184. 10.1038/s41467-018-04580-3.

24 Abraham RT, Weiss A. Jurkat T cells and development of the T-cell receptor signalling paradigm. Nat Rev Immunol 2004;4:301–8. 10.1038/nri1330.

25 Larange A, Takazawa I, Kakugawa K, Thiault N, Ngoi S, Olive ME, et al. A regulatory circuit controlled by extranuclear and nuclear retinoic acid receptor α determines T cell activation and function. Immunity 2023. 10.1016/j.immuni.2023.07.017.

26 Abelin JG, Bergstrom EJ, Rivera KD, Taylor HB, Klaeger S, Xu C, et al. Workflow enabling deepscale immunopeptidome, proteome, ubiquitylome, phosphoproteome, and acetylome analyses of sample-limited tissues. Nat Commun 2023;14:1851. 10.1038/s41467-023-37547-0.

27 Krug K, Mertins P, Zhang B, Hornbeck P, Raju R, Ahmad R, et al. A Curated Resource for Phosphosite-specific Signature Analysis. Mol Cell Proteomics 2019;18:576–93. 10.1074/mcp.TIR118.000943.

28 Subramanian A, Tamayo P, Mootha VK, Mukherjee S, Ebert BL, Gillette MA, et al. Gene set enrichment analysis: a knowledge-based approach for interpreting genome-wide expression profiles. Proc Natl Acad Sci U S A 2005;102:15545–50. 10.1073/pnas.0506580102.

29. Richter MF, Zhao KT, Eton E, Lapinaite A, Newby GA, Thuronyi BW, et al. Phage-assisted evolution of an adenine base editor with improved Cas domain compatibility and activity. Nat Biotechnol 2020;38:883–91. 10.1038/s41587-020-0453-z.

30. Huang TP, Newby GA, Liu DR. Precision genome editing using cytosine and adenine base editors in mammalian cells. Nat Protoc 2021;16:1089–128. 10.1038/s41596-020-00450-9.

31 Kluesner MG, Nedveck DA, Lahr WS, Garbe JR, Abrahante JE, Webber BR, et al. EditR: A Method to Quantify Base Editing from Sanger Sequencing. CRISPR J 2018;1:239–50. 10.1089/crispr.2018.0014.

32 Wang H, Kadlecek TA, Au-Yeung BB, Goodfellow HES, Hsu L-Y, Freedman TS, et al. ZAP-70: an essential kinase in T-cell signaling. Cold Spring Harb Perspect Biol 2010;2:a002279. 10.1101/cshperspect.a002279.

33 Goda S, Quale AC, Woods ML, Felthauser A, Shimizu Y. Control of TCR-Mediated Activation of β1 Integrins by the ZAP-70 Tyrosine Kinase Interdomain B Region and the Linker for Activation of T Cells Adapter Protein. The Journal of Immunology 2004;172:5379–87. 10.4049/jimmunol.172.9.5379.

34 Bottini N, Stefanini L, Williams S, Alonso A, Jascur T, Abraham RT, et al. Activation of ZAP-70 through specific dephosphorylation at the inhibitory Tyr-292 by the low molecular weight phosphotyrosine phosphatase (LMPTP). J Biol Chem 2002;277:24220–4. 10.1074/jbc.M202885200.

35 Di Bartolo V, Mège D, Germain V, Pelosi M, Dufour E, Michel F, et al. Tyrosine 319, a newly identified phosphorylation site of ZAP-70, plays a critical role in T cell antigen receptor signaling. J Biol Chem 1999;274:6285–94. 10.1074/jbc.274.10.6285.

36 Jutz S, Leitner J, Schmetterer K, Doel-Perez I, Majdic O, Grabmeier-Pfistershammer K, et al. Assessment of costimulation and coinhibition in a triple parameter T cell reporter line: Simultaneous measurement of NF-κB, NFAT and AP-1. J Immunol Methods 2016;430:10–20. 10.1016/j.jim.2016.01.007.

37. Li J, Lin J, Huang S, Li M, Yu W, Zhao Y, et al. Functional Phosphoproteomics in Cancer Chemoresistance Using CRISPR-Mediated Base Editors. Adv Sci 2022;9:e2200717. 10.1002/advs.202200717.

38 Pihlajamaa P, Kauko O, Sahu B, Kivioja T, Taipale J. A competitive precision CRISPR method to identify the fitness effects of transcription factor binding sites. Nat Biotechnol 2022. 10.1038/s41587-022-01444-6.

39 Takayama K-I, Suzuki T, Tsutsumi S, Fujimura T, Urano T, Takahashi S, et al. RUNX1, an androgen- and EZH2-regulated gene, has differential roles in AR-dependent and -independent prostate cancer. Oncotarget 2015;6:2263–76. 10.18632/oncotarget.2949.

40 Ritter M, Klimiankou M, Klimenkova O, Schambach A, Hoffmann D, Schmidt A, et al. Cooperating, congenital neutropenia-associated Csf3r and Runx1 mutations activate pro-inflammatory signaling and inhibit myeloid differentiation of mouse HSPCs. Ann Hematol 2020;99:2329–38. 10.1007/s00277-020-04194-0.

41 Quesada AE, Montalban-Bravo G, Luthra R, Patel KP, Sasaki K, Bueso-Ramos CE, et al. Clinico-pathologic characteristics and outcomes of the World Health Organization (WHO) provisional entity de novo acute myeloid leukemia with mutated RUNX1. Mod Pathol 2020;33:1678–89. 10.1038/s41379-020-0531-2.

42 Huang K, Liu X, Li Y, Wang Q, Zhou J, Wang Y, et al. Genome-Wide CRISPR-Cas9 Screening Identifies NF-κB/E2F6 Responsible for EGFRvIII-Associated Temozolomide Resistance in Glioblastoma. Adv Sci 2019;6:1900782. 10.1002/advs.201900782.

43 Zhang Q, Sun M, Zhou S, Guo B. Class I HDAC inhibitor mocetinostat induces apoptosis by activation of miR-31 expression and suppression of E2F6. Cell Death Discov 2016;2:16036. 10.1038/cddiscovery.2016.36.

44 Cheng FHC, Lin H-Y, Hwang T-W, Chen Y-C, Huang R-L, Chang C-B, et al. E2F6 functions as a competing endogenous RNA, and transcriptional repressor, to promote ovarian cancer stemness. Cancer Sci 2019;110:1085–95. 10.1111/cas.13920.

45 Johnson JL, Yaron TM, Huntsman EM, Kerelsky A, Song J, Regev A, et al. An atlas of substrate specificities for the human serine/threonine kinome. Nature 2023;613:759–66. 10.1038/s41586-022-05575-3.

46 Wang B, Wang M, Zhang W, Xiao T, Chen C-H, Wu A, et al. Integrative analysis of pooled CRISPR genetic screens using MAGeCKFlute. Nat Protoc 2019;14:756–80. 10.1038/s41596-018-0113-7.

47 Raudvere U, Kolberg L, Kuzmin I, Arak T, Adler P, Peterson H, et al. g:Profiler: a web server for functional enrichment analysis and conversions of gene lists (2019 update). Nucleic Acids Res 2019;47:W191–8. 10.1093/nar/gkz369.

48 Zeeshan Chaudhry M, Borkner L, Berberich-Siebelt F, Cicin-Sain L. NFAT signaling is indispensable for persistent memory responses of MCMV-specific CD8^+^T cells. bioRxiv 2023. 10.1101/2023.05.02.539029.

49 Feske S. Calcium signalling in lymphocyte activation and disease. Nat Rev Immunol 2007;7:690–702. 10.1038/nri2152.

50 Hogan PG, Chen L, Nardone J, Rao A. Transcriptional regulation by calcium, calcineurin, and NFAT. Genes Dev 2003;17:2205–32. 10.1101/gad.1102703.

51 Hogan PG, Lewis RS, Rao A. Molecular basis of calcium signaling in lymphocytes: STIM and ORAI. Annu Rev Immunol 2010;28:491–533. 10.1146/annurev.immunol.021908.132550.

52 Ortega-Pérez I, Cano E, Were F, Villar M, Vázquez J, Redondo JM. c-Jun N-terminal kinase (JNK) positively regulates NFATc2 transactivation through phosphorylation within the N-terminal regulatory domain. J Biol Chem 2005;280:20867–78. 10.1074/jbc.M501898200.

53 Dejmek J, Säfholm A, Kamp Nielsen C, Andersson T, Leandersson K. Wnt-5a/Ca2+-induced NFAT activity is counteracted by Wnt-5a/Yes-Cdc42-casein kinase 1alpha signaling in human mammary epithelial cells. Mol Cell Biol 2006;26:6024–36. 10.1128/MCB.02354-05.

54 Ishitani T, Kishida S, Hyodo-Miura J, Ueno N, Yasuda J, Waterman M, et al. The TAK1-NLK mitogen-activated protein kinase cascade functions in the Wnt-5a/Ca(2+) pathway to antagonize Wnt/beta-catenin signaling. Mol Cell Biol 2003;23:131–9. 10.1128/MCB.23.1.131-139.2003.

55 MacDonnell SM, Weisser-Thomas J, Kubo H, Hanscome M, Liu Q, Jaleel N, et al. CaMKII negatively regulates calcineurin-NFAT signaling in cardiac myocytes. Circ Res 2009;105:316–25. 10.1161/CIRCRESAHA.109.194035.

56 Anshabo AT, Milne R, Wang S, Albrecht H. CDK9: A Comprehensive Review of Its Biology, and Its Role as a Potential Target for Anti-Cancer Agents. Front Oncol 2021;11:678559. 10.3389/fonc.2021.678559.

57 Phee H, Au-Yeung BB, Pryshchep O, O’Hagan KL, Fairbairn SG, Radu M, et al. Pak2 is required for actin cytoskeleton remodeling, TCR signaling, and normal thymocyte development and maturation. Elife 2014;3:e02270. 10.7554/eLife.02270.

58 Pareek TK, Lam E, Zheng X, Askew D, Kulkarni AB, Chance MR, et al. Cyclin-dependent kinase 5 activity is required for T cell activation and induction of experimental autoimmune encephalomyelitis. J Exp Med 2010;207:2507–19. 10.1084/jem.20100876.

59 Askew D, Pareek TK, Eid S, Ganguly S, Tyler M, Huang AY, et al. Cyclin-dependent kinase 5 activity is required for allogeneic T-cell responses after hematopoietic cell transplantation in mice. Blood 2017;129:246–56. 10.1182/blood-2016-05-702738.

60 Chen M, Pratt CP, Zeeman ME, Schultz N, Taylor BS, O’Neill A, et al. Identification of PHLPP1 as a tumor suppressor reveals the role of feedback activation in PTEN-mutant prostate cancer progression. Cancer Cell 2011;20:173–86. 10.1016/j.ccr.2011.07.013.

61 Nitsche C, Edderkaoui M, Moore RM, Eibl G, Kasahara N, Treger J, et al. The phosphatase PHLPP1 regulates Akt2, promotes pancreatic cancer cell death, and inhibits tumor formation. Gastroenterology 2012;142:377–87.e1–5. 10.1053/j.gastro.2011.10.026.

62 Brognard J, Newton AC. PHLiPPing the switch on Akt and protein kinase C signaling. Trends Endocrinol Metab 2008;19:223–30. 10.1016/j.tem.2008.04.001.

63 Cohen Katsenelson K, Stender JD, Kawashima AT, Lordén G, Uchiyama S, Nizet V, et al. PHLPP1 counter-regulates STAT1-mediated inflammatory signaling. Elife 2019;8.: 10.7554/eLife.48609.

64 Patterson SJ, Han JM, Garcia R, Assi K, Gao T, O’Neill A, et al. Cutting edge: PHLPP regulates the development, function, and molecular signaling pathways of regulatory T cells. J Immunol 2011;186:5533–7. 10.4049/jimmunol.1002126.

65 Schmiedel BJ, Singh D, Madrigal A, Valdovino-Gonzalez AG, White BM, Zapardiel-Gonzalo J, et al. Impact of Genetic Polymorphisms on Human Immune Cell Gene Expression. Cell 2018;175:1701–15.e16. 10.1016/j.cell.2018.10.022.

66 Liu N, Guo Y, Ning S, Duan M. Phosphorylation regulates the binding of intrinsically disordered proteins via a flexible conformation selection mechanism. Communications Chemistry 2020;3:1–9. 10.1038/s42004-020-00370-5.

67 Nicolaou ST, Hebditch M, Jonathan OJ, Verma CS, Warwicker J. PhosIDP: a web tool to visualize the location of phosphorylation sites in disordered regions. Sci Rep 2021;11:9930. 10.1038/s41598-021-88992-0.

68 Trinidad JC, Barkan DT, Gulledge BF, Thalhammer A, Sali A, Schoepfer R, et al. Global identification and characterization of both O-GlcNAcylation and phosphorylation at the murine synapse. Mol Cell Proteomics 2012;11:215–29. 10.1074/mcp.O112.018366.

69 Sharma S, Findlay GM, Bandukwala HS, Oberdoerffer S, Baust B, Li Z, et al. Dephosphorylation of the nuclear factor of activated T cells (NFAT) transcription factor is regulated by an RNA-protein scaffold complex. Proc Natl Acad Sci U S A 2011;108:11381–6. 10.1073/pnas.1019711108.

70 Gwack Y, Sharma S, Nardone J, Tanasa B, Iuga A, Srikanth S, et al. A genome-wide Drosophila RNAi screen identifies DYRK-family kinases as regulators of NFAT. Nature 2006;441:646–50. 10.1038/nature04631.

71 Liu H, Wang K, Chen S, Sun Q, Zhang Y, Chen L, et al. NFATc1 phosphorylation by DYRK1A increases its protein stability. PLoS One 2017;12:e0172985. 10.1371/journal.pone.0172985.

72 Khor B, Gagnon JD, Goel G, Roche MI, Conway KL, Tran K, et al. The kinase DYRK1A reciprocally regulates the differentiation of Th17 and regulatory T cells. Elife 2015;4.: 10.7554/eLife.05920.

73 Thompson BJ, Bhansali R, Diebold L, Cook DE, Stolzenburg L, Casagrande A-S, et al. DYRK1A controls the transition from proliferation to quiescence during lymphoid development by destabilizing Cyclin D3. J Exp Med 2015;212:953–70. 10.1084/jem.20150002.

74 Cruz-Guilloty F, Pipkin ME, Djuretic IM, Levanon D, Lotem J, Lichtenheld MG, et al. Runx3 and T-box proteins cooperate to establish the transcriptional program of effector CTLs. J Exp Med 2009;206:51–9. 10.1084/jem.20081242.

75 Seo H, Chen J, González-Avalos E, Samaniego-Castruita D, Das A, Wang YH, et al. TOX and TOX2 transcription factors cooperate with NR4A transcription factors to impose CD8+ T cell exhaustion. Proc Natl Acad Sci U S A 2019;116:12410–5. 10.1073/pnas.1905675116.

76 Chen J, López-Moyado IF, Seo H, Lio C-WJ, Hempleman LJ, Sekiya T, et al. NR4A transcription factors limit CAR T cell function in solid tumours. Nature 2019;567:530–4. 10.1038/s41586-019-0985-x.

77 Viéitez C, Busby BP, Ochoa D, Mateus A, Memon D, Galardini M, et al. High-throughput functional characterization of protein phosphorylation sites in yeast. Nat Biotechnol 2022;40:382–90. 10.1038/s41587-021-01051-x.

78 Beltrao P, Albanèse V, Kenner LR, Swaney DL, Burlingame A, Villén J, et al. Systematic functional prioritization of protein posttranslational modifications. Cell 2012;150:413–25. 10.1016/j.cell.2012.05.036.

79 Ochoa D, Jarnuczak AF, Viéitez C, Gehre M, Soucheray M, Mateus A, et al. The functional landscape of the human phosphoproteome. Nat Biotechnol 2020;38:365–73. 10.1038/s41587-019-0344-3.

80 Beltrao P, Bork P, Krogan NJ, van Noort V. Evolution and functional cross-talk of protein post-translational modifications. Mol Syst Biol 2013;9:714. 10.1002/msb.201304521.

81 Abelin JG, Bergstrom EJ, Taylor HB, Rivera KD, Klaeger S, Olive ME, et al. MONTE enables serial immunopeptidome, ubiquitylome, proteome, phosphoproteome, acetylome analyses of sample-limited tissues. bioRxiv 2021:2021.06.22.449417. 10.1101/2021.06.22.449417.

82 Burt RA, Dejanovic B, Peckham HJ, Lee KA, Li X, Ounadjela JR, et al. Novel Antibodies for the Simple and Efficient Enrichment of Native O-GlcNAc Modified Peptides. Mol Cell Proteomics 2021;20:100167. 10.1016/j.mcpro.2021.100167.

83 Li H, Remsberg JR, Won SJ, Zhao KT, Huang TP, Lu B, et al. Assigning functionality to cysteines by base editing of cancer dependency genes. bioRxiv 2022:2022.11.17.516964. 10.1101/2022.11.17.516964.

84 Mahrus S, Trinidad JC, Barkan DT, Sali A, Burlingame AL, Wells JA. Global sequencing of proteolytic cleavage sites in apoptosis by specific labeling of protein N termini. Cell 2008;134:866–76. 10.1016/j.cell.2008.08.012.

85 Hwang G-H, Park J, Lim K, Kim S, Yu J, Yu E, et al. Web-based design and analysis tools for CRISPR base editing. BMC Bioinformatics 2018;19:542. 10.1186/s12859-018-2585-4.

86 Chen PJ, Hussmann JA, Yan J, Knipping F, Ravisankar P, Chen P-F, et al. Enhanced prime editing systems by manipulating cellular determinants of editing outcomes. Cell 2021;184:5635–52.e29. 10.1016/j.cell.2021.09.018.

87 Mari T, Mösbauer K, Wyler E, Landthaler M, Drosten C, Selbach M. In Vitro Kinase-to-Phosphosite Database (iKiP-DB) Predicts Kinase Activity in Phosphoproteomic Datasets. J Proteome Res 2022;21:1575–87. 10.1021/acs.jproteome.2c00198.

88 Litichevskiy L, Peckner R, Abelin JG, Asiedu JK, Creech AL, Davis JF, et al. A Library of Phosphoproteomic and Chromatin Signatures for Characterizing Cellular Responses to Drug Perturbations. Cell Syst 2018;6:424–43.e7. 10.1016/j.cels.2018.03.012.

